# Collagen XVIII promotes breast cancer through EGFR/ErbB signaling and its ablation improves the efficacy of ErbB-targeting inhibitors

**DOI:** 10.1101/2022.01.10.474416

**Authors:** Raman Devarajan, Hellevi Peltoketo, Valerio Izzi, Heli Ruotsalainen, Saila Kauppila, Marja-Riitta Väisänen, Gunilla Rask, Guillermo Martínez-Nieto, Sanna-Maria Karppinen, Timo Väisänen, Inderjeet Kaur, Jussi Koivunen, Takako Sasaki, Robert Winqvist, Fredrik Wärnberg, Malin Sund, Taina Pihlajaniemi, Ritva Heljasvaara

## Abstract

The tumor extracellular matrix (ECM) is a critical regulator of cancer progression and metastasis, significantly affecting the treatment response. Expression of collagen XVIII (ColXVIII), a ubiquitous component of basement membranes, is induced in many solid tumors, but its involvement in tumorigenesis has remained elusive. We show here that ColXVIII is markedly upregulated in human breast cancer (BC) cells and is closely associated with a poor prognosis in high-grade BC, especially in human epidermal growth factor receptor 2 (HER2)-positive and basal/triple-negative cases. We identified a novel mechanism of action for ColXVIII as a modulator of epidermal growth factor receptor (EGFR/ErbB) signaling and show that it forms a complex with EGFR, HER2 and α6 integrin to promote cancer cell proliferation in a pathway involving its N-terminal portion and the MAPK/ERK1/2 and PI3K/Akt cascades. *In vivo* studies with *Col18a1* mouse models crossed with the MMTV-PyMT mammary carcinogenesis model showed that the short ColXVIII isoform promotes BC growth and metastasis in a tumor cell-autonomous manner. Moreover, the number of mammary cancer stem cells was significantly reduced in both mouse and human cell models upon ColXVIII inhibition. Finally, ablation of ColXVIII in human BC cells and the MMTV-PyMT model substantially improved the efficacy of certain EGFR/ERbB-targeting therapies, even abolishing resistance to EGFR/ErbB inhibitors in some cell lines. In summary, a new function is revealed for ColXVIII in sustaining the stemness properties of BC cells, and tumor progression and metastasis through EGFR/ErbB signaling, suggesting that targeting ColXVIII in the tumor milieu may have significant therapeutic potential.

**One Sentence Summary:** Collagen XVIII is upregulated in breast cancer and promotes mammary carcinogenesis through EGFR/ErbB signaling and by sustaining cancer stem cells, so that its targeting improves the efficacy of ErbB-targeted therapies.

## INTRODUCTION

Breast cancer (BC) is the most common cancer among women, with over two million new cases diagnosed in 2020, and accounts for 25% of all female cancers *(1)*. Treatment options depend on the type and course of the disease, and on hormone and human growth factor receptor 2 (HER2) status, mutations, proliferation index and differentiation score *(2)*. Hence, BC patients are treated with different combinations of surgery, radiation, chemotherapy and endocrine therapy, as well as with targeted immuno- or small molecule therapies. Despite significant advances in BC care, over 0.6 million women die of BC annually, it accounts for 7% of all female cancer deaths, and the 5-year recurrence rate for all BC cases is around 10% *(1, 3)*.

A major challenge in cancer treatment is intrinsic or acquired drug resistance, which is responsible for most of the relapses that occur after an initially favorable response to treatment *(4, 5)*. For example, approximately 70% of advanced BCs overexpressing the human epidermal growth factor receptor 2 (HER2) develop resistance to trastuzumab, a monoclonal antibody (mAB) targeting HER2, and progress to metastatic disease. Many patients also become resistant to lapatinib, a small-molecule tyrosine kinase inhibitor of HER2, and to epidermal growth factor receptor 1 (EGFR) *(6, 7)*. In addition, residual disease in the breast or lymph nodes after neoadjuvant chemotherapy may carry a high risk of recurrence in patients who present with early-stage triple-negative BC (TNBC), and no targeted molecular therapies are available for this subtype *(8, 9)*.

While genetic alterations in cells predispose to, initiate and drive malignancy, cancer progression is enabled by a dysregulated tumor microenvironment (TME), comprising different types of stromal cells together with the extracellular matrix (ECM) *(10, 11)*. Both tumor and stromal cells actively produce ECM proteins and ECM-modifying enzymes to remodel the TME, which then promotes the growth of cancer cells and their invasion into the surrounding tissue and beyond *(12, 13)*. Moreover, biological and mechanical cues from the ECM support the acquisition of cancer stem cell (CSC) properties by somatic tumor cells, thus favoring continuous tumor growth and the development of drug resistances and eventually disease relapse *(14, 15)*. Our recent analysis has shown that the expression of a variety of ECM components in cancers is precisely regulated by specific oncogenic drivers and downstream transcription factors and correlates with the patient’s prognosis *(16)*. This study and a number of others, including our recent works *(17–19)*, highlight the utility of ECM molecules as diagnostic biomarkers and in disease follow-up and unveil new therapeutic possibilities for inhibiting cancer progression and metastasis and dismantling resistance to cancer therapies by targeting the ECM *(12–15)*.

Collagen XVIII (ColXVIII) is a ubiquitous component of epithelial and endothelial basement membranes (BM) *(20)*. It is a structurally complex and functionally versatile molecule with roles in the eye, nervous system and adipose tissue, for example. ColXVIII exists in three isoforms, short, medium and long, which differ in their N-terminal non-collagenous (NC) domain structure, tissue specificity and functions. All three isoforms contain an endostatin domain, a widely studied BM-derived anti-angiogenic molecule *(20–22)*, in their C-terminal NC1 portion and a laminin-G/thrombospondin-1-like (TSP-1) domain in their N-terminal NC11 portion. The long ColXVIII isoform has two additional domains in the N-terminus, a mucin-like domain (MUCL-C18) and a Wnt-binding Frizzled-like domain (FZ-C18) which is spliced out of the medium ColXVIII isoform *(20, 23)*. In several neoplasms, including lung, prostate, skin and gastric cancers, both ColXVIII overexpression in tumor tissues and high endostatin levels in the patients’ sera have been associated with disease progression and poor prognosis rather than with tumor repression by endostatin *(22)*. However, the mechanism by which ColXVIII promotes tumor growth and progression are still unclear.

We set out here to investigate the role of ColXVIII in BC and its mechanisms of action using genetic mouse tumor models and human BC cell models, and to assess the translational value of ColXVIII by correlating its expression with the clinicopathological features of human BC and by conducting drug tests in ColXVIII-deficient cell and mouse models. Our studies revealed a previously unidentified mechanism for ColXVIII in the regulation of EGFR/ErbB signaling in BC that leads to tumor promotion and demonstrated significant upregulation of ColXVIII expression in high-grade BC, which was associated with a poor clinical outcome. In addition, our preclinical assays showed that ColXVIII targeting has a promising therapeutic potential in the treatment of BC.

## RESULTS

### ColXVIII expression in human breast cancers

Immunohistochemical (IHC) analysis performed on 116 human BC specimens (Table S1) with a monoclonal custom-made ColXVIII antibody (DB144-N2) (Table S2) showed that the ColXVIII signal is prominent in the BMs of blood vessels, mammary ducts and lobules in normal breast tissue adjacent to tumor regions (Fig. 1A, Fig. S1). In addition, ColXVIII can be detected in the thin BM surrounding the adipocytes. In ductal carcinoma *in situ* (DCIS) the ducts filled with tumor cells are usually surrounded by an intact ColXVIII-positive BM/myoepithelial cell layer, albeit the ColXVIII signal may be discontinuous or even completely lacking at some tumor borders (Fig. 1B-C, Fig. S1). Intriguingly, a cytoplasmic ColXVIII staining ranging from weak to moderate can frequently be detected in tumor cells in DCIS (Fig. 1B-C, Fig. S1). In invasive ductal carcinomas (IDC) of various grades ColXVIII expression is commonly seen in the cytoplasm of tumor cells, though the staining intensity varies considerably from weak to strong between samples and tumor regions, being more intense in high-grade tumors (Fig. 1D-F, Fig. S1). In IDCs, ColXVIII expression is either fragmented or completely lost from epithelial BM/myoepithelium around the tumors (Fig. 1D-F). The ColXVIII signal is prominent in the vascular BMs of all DCIS and IDC samples (Fig. 1A-I, Fig. S1), and occasionally also in other stromal cells, including myofibroblasts (Fig. S1). The authenticity of the ColXVIII staining patterns was verified here in several tumor samples with a polyclonal custom-made human ColXVIII antibody (QH48.18) (Fig. S1).

**Fig. 1.**
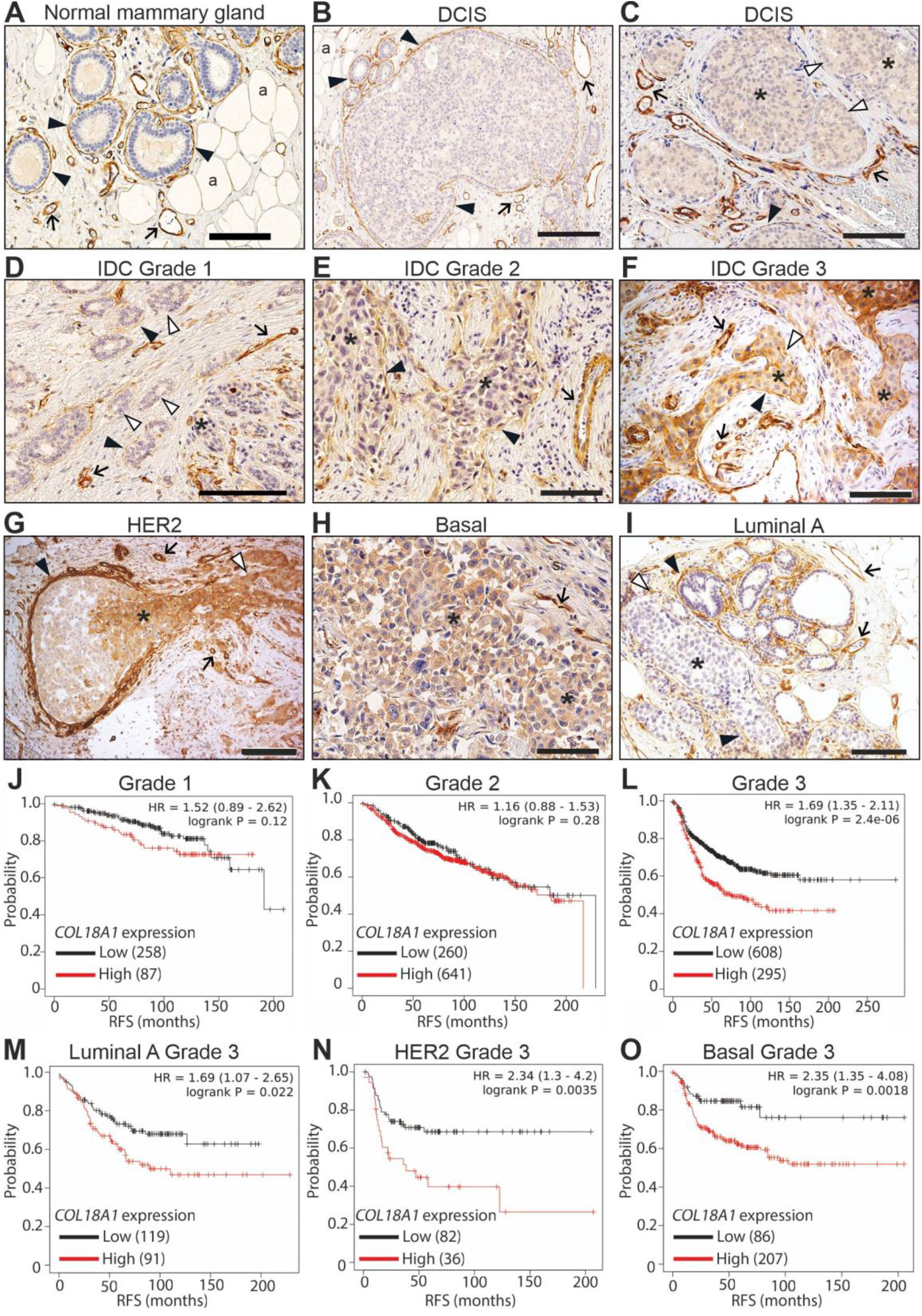
High ColXVIII expression is associated with poor prognosis for human BC. **(A-I)** A monoclonal anti-ColXVIII antibody DB144-N2 was used to detect ColXVIII in the human BC specimens. Representative images of ColXVIII expression and localization in **(A)** normal breast tissue, **(B-D)** ductal carcinoma *in situ* (DCIS), **(D-F)** invasive ductal carcinoma (IDC) of grades 1-3, and the **(G)** HER2, **(H)** basal/TNBC and **(I)** luminal A type of BC. The negative staining control for the DB144-N2 is shown in Fig. S1. Black arrowhead, epithelial basement membrane (BM); white arrowhead, ColXVIII absent in the epithelial BM; arrow, vascular BM; asterisk, cytoplasmic staining in tumor cells; a, adipocyte. Scale bars 100μm. **(J-O)** Kaplan-Meir plots showing the relapse-free survival (RFS) of BC patients stratified by ColXVIII expression levels (probe: 209082_s_at), by cancer grade **(J-L)** and by cancer subtype **(M-O)**. High ColXVIII, red line; low ColXVIII, black line. The open access gene expression data and patients’ survival information from TCGA, GEO and EGA, compiled in a single database at www.kmplot.com *(24)* were used for the meta-analyses. Hazard ratios (HR) and log-rank P values were automatically computed using the best-performing threshold as the cutoff. The initial number of patients in each group (N) is indicated in the survival graphs.

Interesting differences in ColXVIII expression were observed when BC samples were classified according to their molecular subtypes. The cytoplasmic ColXVIII signal is usually strong or moderate in HER2-positive and basal/TNBC cases, and the staining intensity in the BM/myoepithelial cell layer around the tumors varies from negative to strong (Fig. 1G, H). In one case we observed that ColXVIII was markedly upregulated in the cytoplasm of invasive tumor cells but absent from the BM/myoepithelium of the invasive tumor area, whereas the cytoplasmic ColXVIII signal at the DCIS site was weak in spite of the fact that both the BM/myoepithelium and the endothelium showed strong ColXVIII staining (Fig. 1G). The ColXVIII signals in the samples of the luminal A subtype were variable both in the cytoplasm and around the tumor nests, ranging from negligible to prominent staining (Fig. 1I).

Analyses of open databases that include survival data on BC patients to estimate the association between ColXVIII mRNA levels and patient survival *(24)* showed that high ColXVIII expression was significantly associated with poor prognosis in patients with high-grade BCs, where the hazard ratio exceeded 1.5, but not in unclassified patients or in those with low-grade tumors (Fig. 1J-L, Fig. S2). When the patients with high-risk grade 3 cancers were further categorized into major molecular subtypes, high ColXVIII was more significantly associated with poor survival in the HER2, basal/TNBC and luminal B subgroups than in the luminal A subgroup (Fig.1 M-O, Fig. S3), indicating that ColXVIII upregulation correlates with recurrence in the case of high-grade BCs.

An in-house indirect ELISA assay was performed to quantify the plasma levels of N-terminal ColXVIII fragments in a small number of healthy controls and BC patients. Most of the patient samples showed higher plasma ColXVIII levels than the healthy controls, and the average plasma ColXVIII concentration in HER2, and particularly in the basal/TNBC subtypes, but not in luminal A cases, was significantly higher than in the controls (Fig. S4A). When the same data were grouped according to metastatic status, plasma ColXVIII levels were significantly higher in lymph node-positive than in node-negative luminal A cases, suggesting that ColXVIII could predict tumor metastasis in this BC subtype (Fig. S4B). On the other hand, ColXVIII levels in the plasma of HER2 and basal/TNBC-type patients were high in both node-negative and node-positive cases, implying that ColXVIII could be used as an early diagnostic marker for these subtypes, even before metastasis (Fig. S4B).

### ColXVIII promotes tumor cell proliferation through its N-terminal TSP-1 domain

ColXVIII signals in cultured human BC cell lines were prominent in the HER2-amplified JIMT-1 cells, the triple-negative MDA-MB-231 and HS578T cells and the MCF7 cells representing the luminal A subtype (Fig. 2A-B). In the other BC cell lines tested, including T47D (luminal A), BT474 (luminal B) and SKBR3 (HER2), and in the non-cancerous breast epithelial cell line MCF10A, ColXVIII was also present but showed some variation (Fig. 2A-B).

**Fig. 2.**
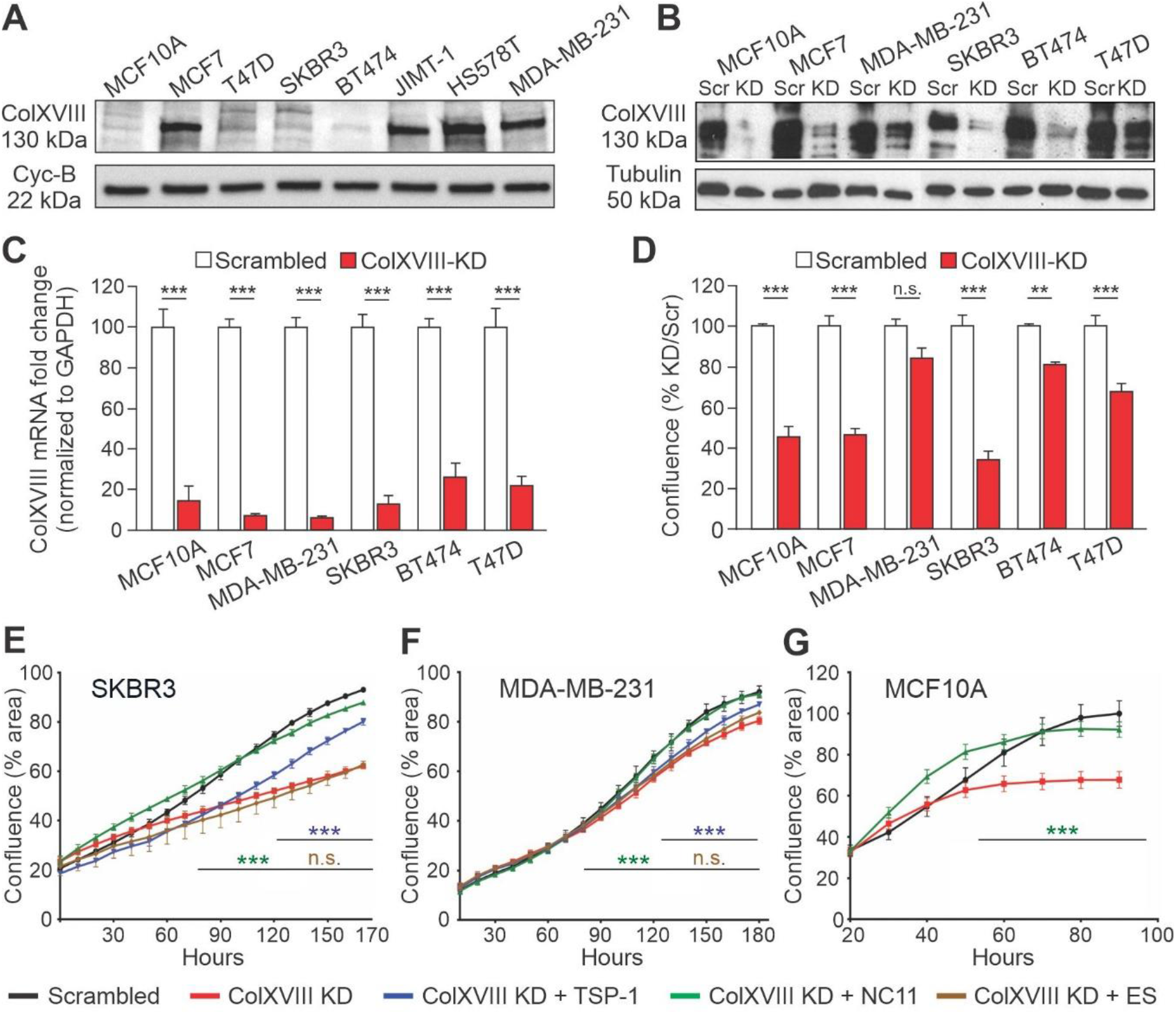
ColXVIII promotes BC cell proliferation through its N-terminal domain. **(A)** A representative immunoblot of ColXVIII expression in human BC and mammary epithelial cell lysates. The size of the major ColXVIII band, ∼130 kDa, corresponds to the core polypeptide of the short ColXVIII isoform. The loading control is cyclophilin B. **(B)** A representative immunoblot of ColXVIII protein levels in various ColXVIII knockdown (KD) cell lines and corresponding scrambled controls (Scr). The loading control is β−tubulin. **(A, B)** A polyclonal ColXVIII antibody, QH48.18, was used in the immunoblots. **(C)** RT-qPCR analysis of ColXVIII mRNA levels after KD in the cell lines indicated. **(D)** Confluency of ColXVIII KD cell cultures relative to Scr cultures (%) of the cell lines indicated, measured by an IncuCyte live cell analysis system for 96 hours. **(E-G)** The graphs indicate the cell confluency measured upon the addition of recombinant non-collagenous (NC) ColXVIII fragments (500 ng/ml) to the KD cells. TSP-1, N-terminal laminin-G/thrombospondin-1-like domain; NC11, full-length N-terminal NC11 fragment containing TSP-1-like MUCL-18 and FZ-C18 domains; ES, endostatin domain. The data in C-G are presented as means ± s.e.m. The two-tailed Student’s ‘t’ test was used in the statistical analyses (treated vs. Scr); **P<0.01, ***P<0.001, n.s., not significant. n=6-7 technical replicates in C and n=3 technical replicates in D-G.

To investigate the role of ColXVIII in breast carcinogenesis, we inhibited its expression in human BC cell lines with RNA interference. A mixture of two small interfering RNAs (siRNA), one targeting the TSP-1 region at the N-terminus of ColXVIII and the other targeting the C-terminal endostatin, was used to achieve an efficient knockdown (KD) of ColXVIII in selected BC cell lines and in MCF10A cells. Typically, a 70-90% inhibition in mRNA synthesis was achieved by this approach as compared with the scrambled vector transfected control cells, leading to a reduced amount of ColXVIII protein (Fig. 2B-C). Except for the MDA-MB-231 cells, ColXVIII KD significantly reduced the proliferation of human BC cells, ranging from an approximately 20% reduction in the BT474 cells to an almost 70% reduction in the SKBR3 cells during a 96-hour follow-up period (Fig. 2D).

To confirm that the reduction in cell proliferation was caused by ColXVIII KD, and to determine which portion of the ColXVIII molecule could convey this effect, recombinant fragments of various NC domains of ColXVIII were added to SKBR3 KD and MDA-MB-231 KD cells and cell proliferation was recorded for 180 hours. Both the TSP-1 fragment and the full-length N-terminal NC11 fragment (containing TSP-1, MUCL-18 and FZ-C18) were able to reverse the inhibitory effect of siRNA-mediated ColXVIII depletion and restore the proliferation activity of the KD cells to the level of scrambled cells, especially in the HER2-positive SKBR3 cell line (Fig. 2E). By contrast, recombinant endostatin could not rescue the reduced proliferation of SKBR3 KD cells. In the MDA-MB-231 cells the effects of both ColXVIII KD and added N-terminal TSP-1 and NC11 fragments were less impressive, but nevertheless statistically significant, whereas endostatin did not affect the KD cells (Fig. 2F). The non-cancerous MCF10A KD cells also regained their proliferation activity upon the addition of an exogenous NC11 fragment (Fig. 2G). Hence, the results of these *in vitro* experiments suggest that specifically the N-terminal portion of ColXVIII, and even the TSP-1 domain alone, can constitute an ECM signal that activates BC and mammary epithelial cell proliferation.

### ColXVIII supports mammary carcinogenesis in the MMTV-PyMT mouse model

Consistent with the results of human tissue analyses, ColXVIII signals in healthy mouse mammary tissue can be detected around adipocytes, in vascular BMs and in the mammary duct BMs, where it is located next to the alpha smooth muscle actin (αSMA), a marker of myoepithelial cells in mammary ducts and smooth muscle cells in blood vessels (Fig. 3A). As expected, the mammary glands (MG) and adipose tissue of healthy *Col18a1*^−/−^ mice are not reactive with the anti-ColXVIII antibody, whereas the αSMA signal can be observed in the ducts and vessels (Fig. 3A). In MMTV-PyMT (PyMT) mouse mammary tumor tissues the ColXVIII signal is clearly increased and is located around tumor nests and the vascular BMs (Fig. 3B). In addition, a prominent cytoplasmic ColXVIII staining of tumor cells is evident in the late-stage PyMT tumors (Fig. 3B).

**Fig. 3.**
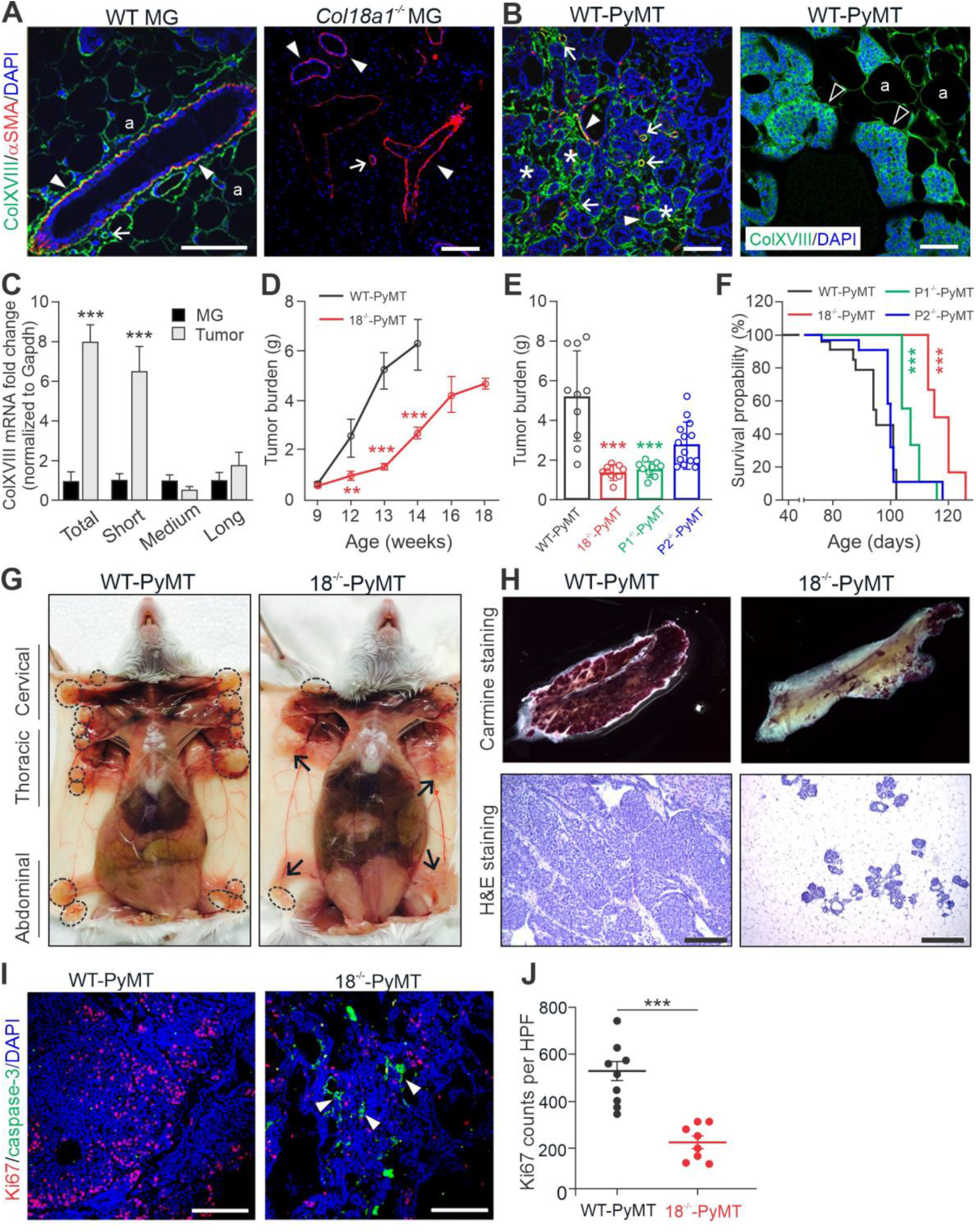
The short ColXVIII isoform promotes mammary tumor growth in mice. **(A-B)** A polyclonal antibody against the N-terminal TSP-1 domain of mouse ColXVIII (green) (Table S2) was used in immunostainings of mammary glands (MG) from healthy wild type (WT) and ColXVIII knockout (*Col18a1*^−/−^) females **(A)** and from wild type MMTV-PyMT (WT-PyMT) mammary tumors **(B)**. Alpha smooth muscle actin (αSMA; red) antibody detects myoepithelial cells in mammary ducts and vascular smooth muscle cells in arteries and veins. Strong cytoplasmic ColXVIII staining of tumor cells can be observed in the late-stage tumors in which the intact BM is sparse (B, right). Arrowhead, basement membrane (BM); arrow, vascular BM; asterisk, tumor nests; a, adipocyte. Scale bars 100μm in A, 200μm in B left, and 100μm in B right. **(C)** Quantification of the three ColXVIII mRNA transcripts (short, medium and long) in WT-PyMT tumor tissues (n=3) normalized to *Gapdh*. N-terminal NC sequences were amplified by qRT-PCR with isoform-specific primers and total ColXVIII mRNA with primers from the C-terminal common endostatin sequence (Table S3). **(D)** Total mammary tumor burden in WT-PyMT and in 18^−/−^-PyMT mice lacking all ColXVIII isoforms at 9-18 weeks of age. At least three WT-PyMT and 18^−/−^-PyMT mice are included at each time point. **(E)** Mammary tumor burden at week 13 in WT-PyMT (N=10) and 18^−/−^-PyMT mice (N=9), and in P1-PyMT mice lacking specifically the short ColXVIII (N=9) and P2-PyMT mice lacking the medium and long ColXVIII (N=14). **(F)** Kaplan-Meier survival plots for the WT-PyMT (n=38), 18^−/−^-PyMT (n=31), P1-PyMT (n=28) and P2-PyMT (n=33) mice. **(G)** Representative photographs of mammary tumors in the WT-PyMT and 18^−/−^-PyMT mice at week 13. The dotted circles mark tumors in the cervical, thoracic and abdominal mammary fat pads. Arrows, macroscopically normal MGs in the 18^−/−^-PyMT mice. **(H)** Representative images of Carmine Alum-stained whole mount preparations of MGs and haematoxylin-eosin (H&E)-stained mammary tumor sections of the WT-PyMT and 18^−/−^-PyMT mice at week 13. Scale bar in the H&E images 200μm. **(I)** Ki67 (red) and cleaved caspase-3 (green) immunostainings of mammary tumors of the 13 weeks-old WT-PyMT and 18^−/−^-PyMT mice. Scale bar 200μm. Arrowheads, clusters of apoptotic cells in the 18^−/−^-PyMT specimen. **(J)** Quantification of the Ki67-positive cell counts in WT-PyMT (n=9) and 18^−/−^-PyMT (n=8) tumors. Four random fields per tumor, imaged with 20x objective, were counted. Statistical analyses: C, D, J: two-tailed Student’s ‘t’ test; E: one-way ANOVA, Bartlett’s post-correction test for equal variances; F: by comparison with WT-PyMT mice in the Mantel-Cox test; **P<0.01; ***P<0.001. Error bars indicate s.e.m.

A qRT-PCR analysis showed that the short ColXVIII isoform in particular is upregulated approximately 7-8-fold in PyMT tumors by comparison with normal mouse mammary tissue (Fig. 3C). To investigate more closely the expression and functions of distinct ColXVIII isoforms in mammary tumors, we established an 18^−/−^-PyMT mouse line lacking all the ColXVIII isoforms, a P1-PyMT line lacking only the short isoform and a P2-PyMT line lacking the medium/long ColXVIII isoforms. Immunostaining of mammary tissues from these crosses confirmed that the ColXVIII signals in the mammary tumors, both around the tumor nests and in the tumor vasculature, result from the short isoform, as the 18^−/−^-PyMT and P1-PyMT tumor tissues did not show any ColXVIII staining whereas the ColXVIII signals from the P2-PyMT tumors were comparable with those from the WT-PyMT specimens (Fig. S5A).

The overall tumor burden was markedly reduced in the18^−/−^-PyMT mice by comparison with the WT-PyMT mice from week 10 onwards, the difference being statistically significant at weeks 12-14 (Fig. 3D-E). In line with the qRT-PCR analysis and immunostainings of PyMT tumor tissues that revealed the involvement of short ColXVIII in mammary carcinogenesis, the tumor burden was approximately 75% lower in both the P1-PyMT and 18^−/−^-PyMT females at week 13 than that in the WT-PyMT females at the same time point (Fig. 3D-E). No further comparison between the WT-PyMT and 18^−/−^-PyMT groups was possible, however, since all the WT-PyMT mice reached the humane end point of the experiment by the age of 14 weeks, whereas the 18^−/−^-PyMT mice could be followed until week 18. The tumor burden in the P2-PyMT mice was slightly lower than in the WT-PyMT group, but the difference was not statistically significant (Fig. 3E). In agreement with these data, the survival rates of the 18^−/−^-PyMT and P1-PyMT mice were significantly better than those of the WT-PyMT and P2-PyMT mice (Fig. 3F).

The tumors in the WT-PyMT mice were found in the cervical, thoracic and abdominal MGs, whereas those in the 18^−/−^-PyMT mice were located mainly in the cervical MGs (Fig. 3G). Whole mount Carmine-Alum and the haematoxylin-eosin stainings showed substantially less cancerous tissue in the MGs of the 13-week-old 18^−/−^-PyMT mice than in those of the WT-PyMT mice, in which the MGs were filled with tumors (Fig. 3H). Moreover, whereas most mammary tumors in the control mice transformed to carcinomas around week 10 and all the WT-PyMT tumors could be classified as carcinomas at week 13, those in the 18^−/−^-PyMT transformed to carcinomas much later, around weeks 16-18, and presented as hyperplasia or adenomas at weeks 8-14 (Fig. 3H, Fig. S5B-C).

There was a significant delay and impaired growth in pulmonary metastasis in the 18^−/−^-PyMT and P1-PyMT mice compared with the WT-PyMT and P2-PyMT mice. Hence all the WT-PyMT and P2-PyMT sacrificed at the age of 13-15 weeks, and 90% and 60%, respectively, of those sacrificed at the age of 10-12 weeks had macrometastases in the lungs. By contrast, the 18^− /−^-PyMT and P1-PyMT mice developed lung metastases at later time points, so that only 10% of 18^−/−^-PyMT mice and 20% of the P1-PyMT mice had lung metastases at weeks 13-15, and 70% and 100%, respectively, at weeks 16-18 (Fig. S6). Overall, out of the mice studied, the WT-PyMT mice had the highest (95%) and the 18^−/−^-PyMT mice had the lowest proportion (27%) of lung metastases during the follow-up period (Fig. S6C). Image analysis revealed significantly larger tumor areas in the lungs of the WT-PyMT and the P2-PyMT mice than in those of the 18^−/−^-PyMT and P1-PyMT mice (Fig. S6D).

Finally, to study the cellular effects of *Col18a1* deletion on mammary carcinogenesis, tumor tissue samples collected from the WT-PyMT and 18^−/−^-PyMT mice were stained for the Ki67 proliferation marker and pro-apoptotic cleaved caspase-3. The average number of Ki67-positive cells was approximately 60% lower in the 18^−/−^-PyMT tumors than in WT-PyMT tumors, indicating that cancer cell proliferation is compromised in the absence of ColXVIII (Fig. 3I-J). Clusters of caspase-3-positive tumor cells were frequently detected in the 18^−/−^-PyMT tumors, whereas in the WT-PyMT tumors their amount was negligible and could not be quantified (Fig. 3I).

### ColXVIII has an autocrine stimulatory function in mammary carcinoma cells

Reciprocal orthotopic allograft transplantation experiments between the WT and *Col18a1*^−/−^ genotypes were performed to determine whether the tumorigenic functions of ColXVIII are tumor cell-autonomous or microenvironmental. Both the WT and *Col18a1*^−/−^ females that received WT-PyMT tumor cells developed palpable tumors by week 7 after implantation (Fig. 4A), but these grew faster in the WT-FVB hosts, reaching an average volume of 550 mm^3^, the humane end point size limit, by week 10. In the *Col18a1*^−/−^*-*FVB hosts the tumors were on average half the size of those in the WT-FVB hosts by the same week, and although the difference was not statistically significant, this observation suggested that host-derived ColXVIII can also contribute to the regulation of tumor growth. As surmised from the previous experiments (Fig. 2 and Fig. 3), cells isolated from the 18^−/−^-PyMT tumors grew much more slowly and developed palpable tumors 5 weeks later, at week 12, in both hosts. Interestingly, the 18^−/−^-PyMT tumors also grew faster in the WT mice than in the *Col18a1*^−/−^ mice, reaching a size of around 400 mm^3^ in 17 weeks whereas those in the *Col18a1*^−/−^ hosts did not exceed 200 mm^3^ in volume within the same time frame (Fig. 4A). This again suggests a contribution of stromal ColXVIII to tumor growth. Ki67 immunostaining of the allografts showed that in both hosts the implanted 18^−/−^-PyMT tumor cells proliferated significantly less than the WT-PyMT cells (Fig. 4B-C). On the other hand, there was no statistical difference in the Ki67 scores for either the 18^−/−^-PyMT cells or the WT-PyMT cells between the hosts (Fig. 4C).

**Fig. 4.**
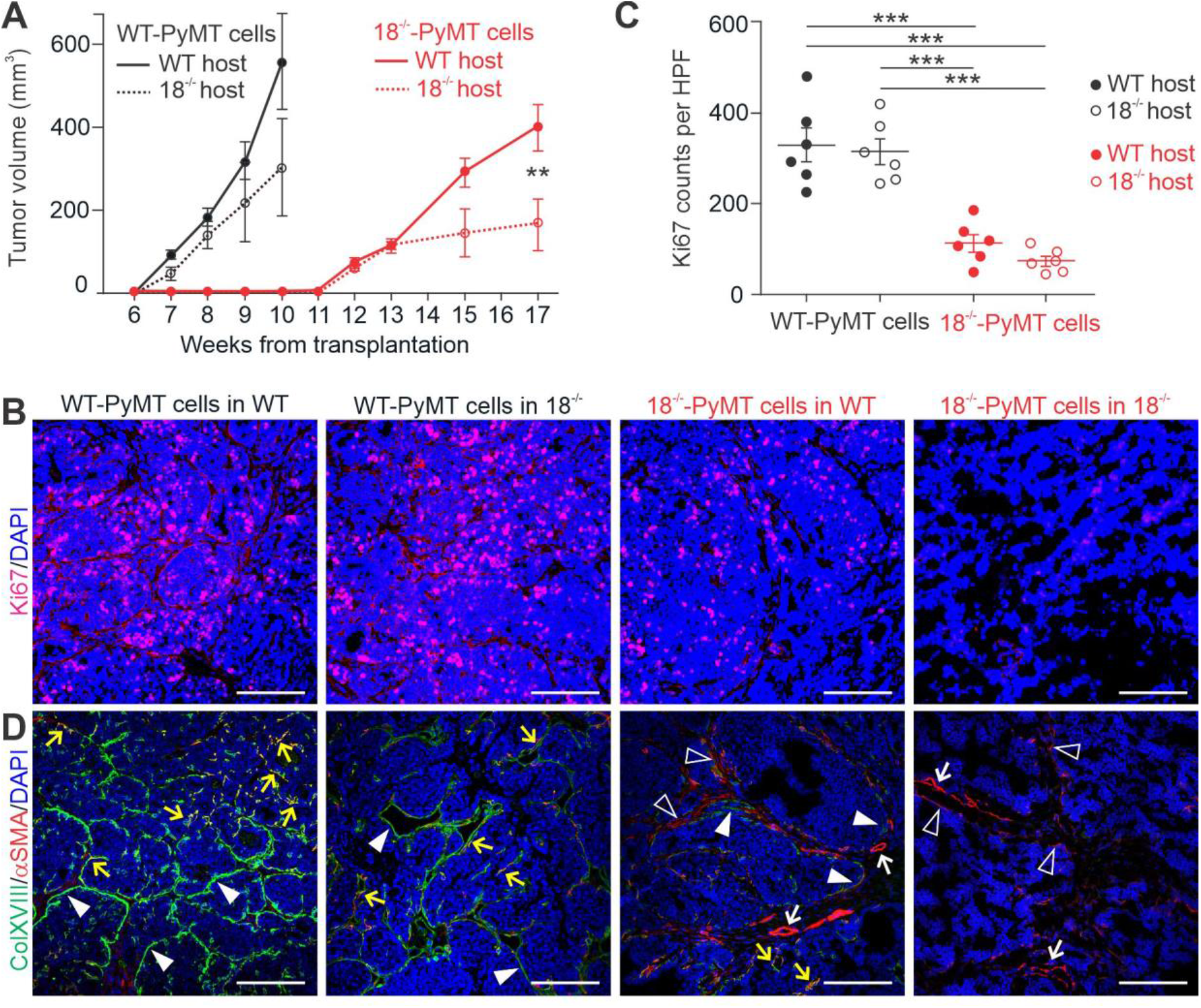
Orthotopic allograft transplantation experiments. **(A)** Growth rates of transplanted WT-PyMT and 18^−/−^-PyMT tumors in WT and *Col18a1*^−/−^ hosts. Numbers of mice (N) and allograft tumors (n): WT-PyMT cells in a WT host and 18^−/−^-PyMT cells in a WT host (N=12, n=24), WT-PyMT cells in *Col18a1*^−/−^ hosts and 18^−/−^-PyMT cells in *Col18a1*^−/−^ hosts (N=6, n=12). **(B)** Tumor cell proliferation. Representative images of Ki67 immunostaining in the WT-PyMT and 18^−/−^-PyMT allografts. **(C)** Quantification of the Ki67-positive cell counts in transplanted tumors. Four random fields per tumor (N=6 per group), imaged with a 20x objective, were counted. The statistical significances of differences in A and C were evaluated using the two-tailed Student’s ‘t’ test. **P<0.01; ***P<0.01. Error bars indicate s.e.m. **(D)** Representative images of ColXVIII (green) and αSMA (red) expression in the allograft tumors. Arrowhead, ColXVIII-positive structures or cells at tumor borders; open arrowhead, αSMA-positive stromal cells; yellow arrow, αSMA and ColXVIII double-positive structures and cells; white arrow, αSMA-positive blood vessels. Scale bars in panels B and D, 200μm.

Immunostaining of the WT-PyMT tumor allografts showed prominent ColXVIII signals at the borders of the tumor nests in the WT hosts but somewhat weaker signals at these sites in the *Col18a1*^−/−^ hosts (Fig. 4D). At some sites the ColXVIII signals overlapped with αSMA present in the myoepithelial/endothelial cell layer of the tumor border. ColXVIII signals were very rare when 18^−/−^-PyMT cells were injected into the WT host, although in some samples faint, discontinuous ColXVIII staining could be observed in the vicinity of αSMA-positive stroma at the tumor borders (Fig. 4D). When 18^−/−^-PyMT cells formed tumors in the *Col18a1*^−/−^ hosts, ColXVIII signals were completely lacking, as expected, and only an αSMA-positive myoepithelium and vasculature could be detected (Fig. 4D).

### ColXVIII supports breast cancer stem cells

High ColXVIII expression has been observed in human mammary stem and progenitor cell populations *(25)* as also in various tissue-specific stem cell niches *(22)*. Using fluorescence-activated cell sorting (FACS), we found that the frequency of mouse mammary CSCs, defined as CD49f^high^ (integrin α6^high^), CD29^high^ (integrin β1^high^), hyaluronan receptor CD44^+^ and heat stable antigen CD24^+^ cell populations *(26)*, was reduced almost by 90% in the 18^−/−^-PyMT tumors by comparison with the WT-PyMT tumors (Fig. 5A). In addition, immunostaining of tumor tissues showed a notable reduction in the integrin β1 signal in the 18^−/−^-PyMT tumors relative to the controls, so that the number of cells that were double positive for integrins β1 and α6 was very low in the knockout tumors (Fig. 5B). Cytokeratin-5 (CK5) is a marker of mature myoepithelial cells when it is co-expressed with αSMA, but discrete CK5^+^αSMA^−^ cells are regarded as CSCs *(27)*. Single positive CK5^+^ cells were abundant inside the WT-PyMT tumor nests, whereas they were approximately 40% less frequent in the 18^−/−^-PyMT tumors (Fig. 5C-D), further confirming that there are less CSCs in knockout tumors. Moreover, immunostaining of allograft tumors showed more CK5^+^ and αSMA^−^ CSCs in WT-PyMT tumors grown in both WT and *Col18a1*^−/−^ hosts, whereas 18^−/−^-PyMT tumors showed more CK5^+^ and αSMA^+^ double-positive cells, as is indicative of reduced CSC characteristics and myoepithelial differentiation in these cells (Fig. 5E).

**Fig. 5.**
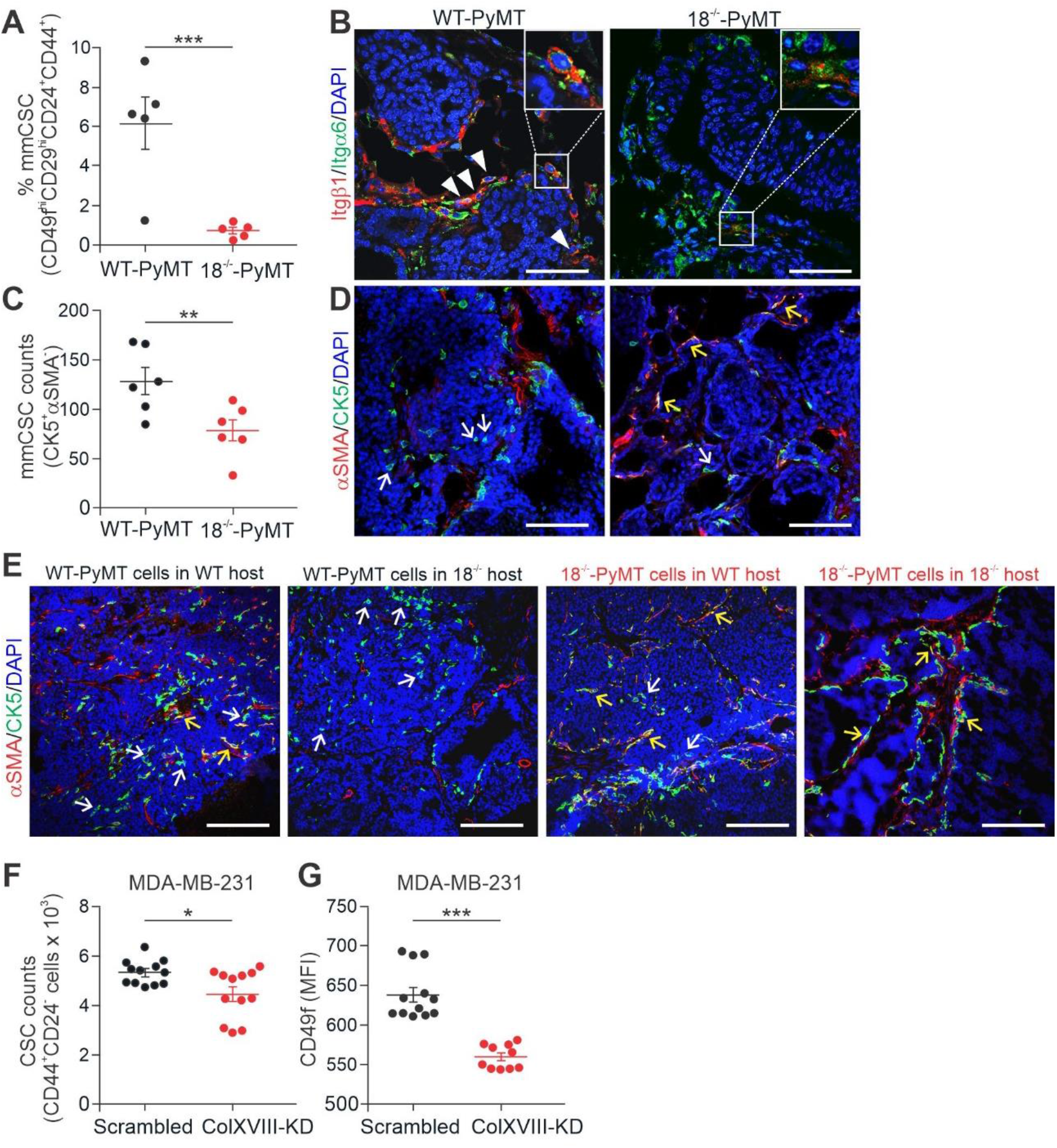
ColXVIII in breast cancer stem cells (CSC). **(A)** Quantification of CD44^+^, CD24^+^, CD29^hi^ and CD49f ^hi^ mouse mammary CSCs sorted by FACS from tumors of the 13-weeks-old WT-PyMT and 18^−/−^-PyMT mice (N=5 per genotype). **(B)** Representative images of CD29 (red) and CD49f (green) immunofluorescence staining of mammary tumors at week 13. Arrowhead, CD29 and CD49f double-positive cells in WT-PyMT. The magnifications in the inserts show strongly double-positive cells in the WT-PyMT tumors but only weakly double-positive ones in the 18^−/−^-PyMT tumors. **(C-D)** Analysis of CK5 and αSMA expression in WT-PyMT and 18^−/−^-PyMT tumor tissues at week 13. (**C)** Quantification of discrete CK5^+^ and αSMA^−^ cells in the tumors. Four random fields per tumor (N=6 per group), imaged with a 20x objective, were counted. **(D)** Representative images of CK5 (green) and αSMA (red) staining. **(E)** Representative images of CK5 (green) and αSMA (red) staining in allograft tumors. White arrows in D and E, CK5-positive and αSMA-negative progenitor cells; yellow arrows, CK5/αSMA double-positive mature myoepithelial cells. **(F)** CSC populations in the ColXVIII siRNA-transfected KD and scrambled vector transfected control MDA-MB-231 cells, as estimated by FACS sorting of the CD44^+^ CD24^low/−^ cells. **(G)** Quantification of the mean fluorescence intensity (MFI) of the CD49f-positive cells in the ColXVIII KD and control MDA-MB-231 cells. Scale bar in B, D and E, 100μm. The statistical significances of the differences in A, C, F and G were evaluated using the two-tailed Student’s ‘t’ test. *, P<0.05; **, P<0.01; ***, P<0.001. Error bars indicate s.e.m.

We then analyzed the CD44^+^ and CD24^low/−^ CSC populations *(28)* in the WT and KD MDA-MB-231 human BC cells using FACS. A significant reduction in the frequency of this CSC population was observed in the siRNA-based ColXVIII KD when compared with the scrambled-treated MDA-MB-231 cells (Fig. 5F), resulting in a remarkable decrease in the mean fluorescence intensity levels of CD49f (Fig. 5G). Moreover, the common stem cell-related transcription factors *NANOG, SNAI1, SNAI2 (SLUG)* and *SOX2 (29)* were downregulated in the MDA-MB-231 KD cells (Fig. S7A), as also in the MCF7 cells with ColXVIII KD (Fig. S7B). Interestingly, ColXVIII expression was significantly higher in the CSC-enriched (CD44^+^CD24^low/−^) subpopulation of MCF7 cells than in the non-CSC subpopulation (CD44^+^CD24^high^) (Fig. S7C). When these subpopulations were assessed in a colony-forming assay, the CSC-enriched MCF7 population formed irregular, heterogenous colonies whereas the non-CSC population formed well-polarized round colonies of approximately equal size (Fig. S7D). Moreover, ColXVIII KD in the non-tumorigenic MCF10A cells reduced the frequency of the CD44^+^CD24^low/−^ population by more than half and led to a significant decrease in the stem cell marker mRNA levels (Fig. S7E-F). Our data thus show that high ColXVIII expression is associated with cellular stemness, and its ablation leads to a significant decrease in the number of tumor-promoting mammary CSC populations.

### ColXVIII is co-expressed with EGFR and HER2 in human breast cancer cells

Our open data analyses have indicated that high ColXVIII expression is associated with poor survival in HER2-amplified and basal/TNBC types of human BC (Fig. 1, Fig. S2, Fig. S3), prompting further investigations into the role of ColXVIII in the EGFR/ErbB signaling pathway. Initial IHC analysis of HER2-type human BC specimens (N=21, Table S1) showed that ColXVIII, EGFR and HER2 are expressed in the same tumor areas that have a high number of Ki67-positive cells (Fig. 6A-B). We then analyzed the expression of ColXVIII and EGFR in a larger BC tissue microarray (TMA) (N=95, Table S1) which had previously been scored for HER2 and Ki67 expression and the nuclear grade. This analysis confirmed that, especially in the HER2 subgroup, strong or moderate cytoplasmic ColXVIII expression in tumor cells correlated with HER2 amplification, and most of these samples also showed strong or moderate EGFR expression (Fig. 6C). In addition, high ColXVIII expression was associated with tumor grade 3 in all the HER2 cases. Correspondingly, a high or moderate cytoplasmic ColXVIII signal was associated with Ki67 expression in almost 60% of the luminal B and TNBC cases and with tumor grades 2 or 3 in 35%, whereas ColXVIII and EGFR were abundantly co-expressed only in a few luminal B specimens in which the tumor cells were also HER2-positive (Fig. 6C). ColXVIII and EGFR signals were found juxtaposed or overlapping in normal human and mouse mammary ducts, with ColXVIII showing a typical BM staining and EGFR signals being localized in the myoepithelial layer (Fig. 6D-E).

**Fig. 6.**
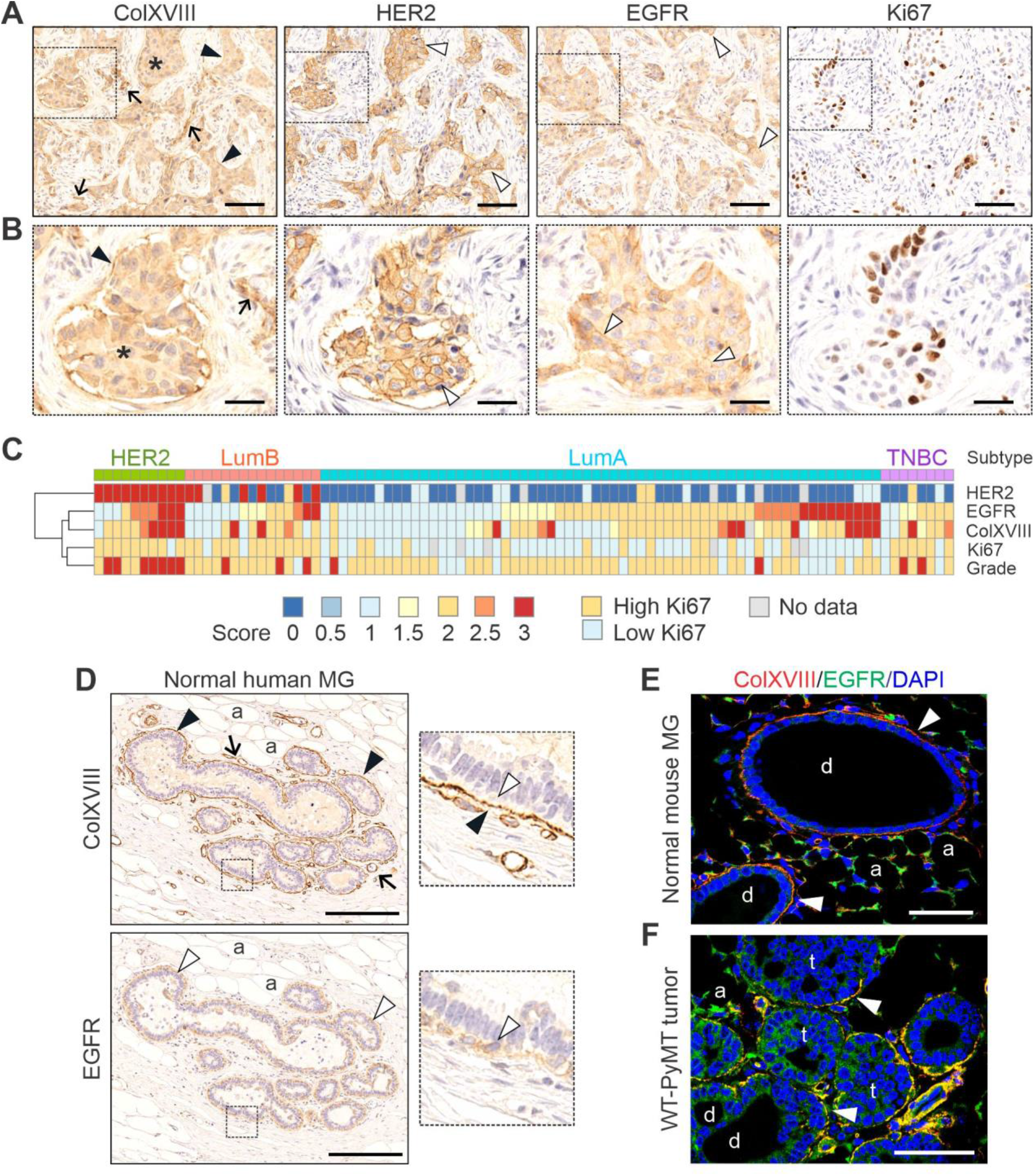
Expression of ColXVIII and EGFR/ErbB in breast tumors and mammary glands. **(A)** Representative images of immunohistochemical staining (IHC) for ColXVIII, EGFR, HER2 and Ki67 in sequential sections of human HER2 type BC. Scale bar, 200μm. Asterisk, cytoplasmic ColXVIII signal in tumor nests; arrow, vascular BM; black arrowhead, epithelial BM; white arrowhead, EGFR or HER2 signal on the plasma membrane. **(B)** Three-fold magnification of the regions indicated in A. Scale bar 600μm. **(C)** Heat map showing IHC scores for HER2, EGFR, ColXVIII and Ki67 and tumor grades, grouped by BC subtypes. Numbers of samples analyzed for each molecular subtype: HER2, N=10; luminal B, N=15; luminal A, N=62; and TNBC, N=8. Scores: 2.5−3 = strong; 1.5−2 = moderate; 0.5−1 = weak; 0 = negative. **(D)** IHC for ColXVIII and EGFR in a normal human mammary gland (MG). Panels on the right depict five-fold magnification of the regions indicated in the left panels and show the juxtaposed localization of ColXVIII and EGFR in the MG epithelial BM and in the myoepithelial cells, respectively. Arrow, vascular BM; black arrowhead, epithelial BM; white arrowhead, myoepithelial cell; a, adipocyte. Scale bar, 100μm. **(E-F)** Immunofluorescence staining for ColXVIII (green) and EGFR (red) in the normal mouse mammary duct and in a wild type MMTV-PyMT (WT-PyMT) mammary tumor. White arrowhead, epithelial BM; a, adipocyte; d, duct; t, tumor nest. Scale bars, 100 μm.

### ColXVIII forms a complex with EGFR and integrin α6 and regulates EGFR/ErbB signaling

Immunofluorescence showed ColXVIII and EGFR co-expression in the basal type MDA-MB-231 (Fig. 7A) and in the HER2-amplified JIMT-1 (Fig. S8A) human BC cells. As cooperation between EGFR and ECM receptor integrins is known to promote the progression and aggressiveness of solid tumors *(15, 30)*, the expression of α6 and β1 integrins, the key integrins in the mammary epithelium, was also analyzed. These integrins are also determinants of breast CSCs, the incidence of which was found to be reduced upon ColXVIII ablation (Fig. 5, Fig. S7). Immunostainings revealed that both the α6 and β1 subunits are expressed with ColXVIII and EGFR in MDA-MB-231 cells (Fig. 7B; Fig. S8B). Proximity ligation assays demonstrated potential interactions between ColXVIII and EGFR in MDA-MB-231 and JIMT-1 cells, as well as between ColXVIII and α6 integrin in MDA-MB-231 cells (Fig. 7C, Fig. S8C). Consistently with this, co-immunoprecipitation assays showed that EGFR and integrin α6 antibodies pull down ColXVIII in MDA-MB-231 and JIMT-1 lysates (Fig. 7D, Fig. S8D-E), and HER2 in JIMT-1 lysates (Fig. 7E). EGFR antibodies also immunoprecipitated HER2 in SKBR3 lysates, and the ColXVIII antibody pulled down both EGFR and HER2 in these cells (Fig. S8F) and EGFR in MCF10A cells (Fig. S8G). Neither EGFR nor ColXVIII antibodies pulled the integrin β1 subunit down in MDA-MB-231 cells (Fig. S8H).

**Fig. 7.**
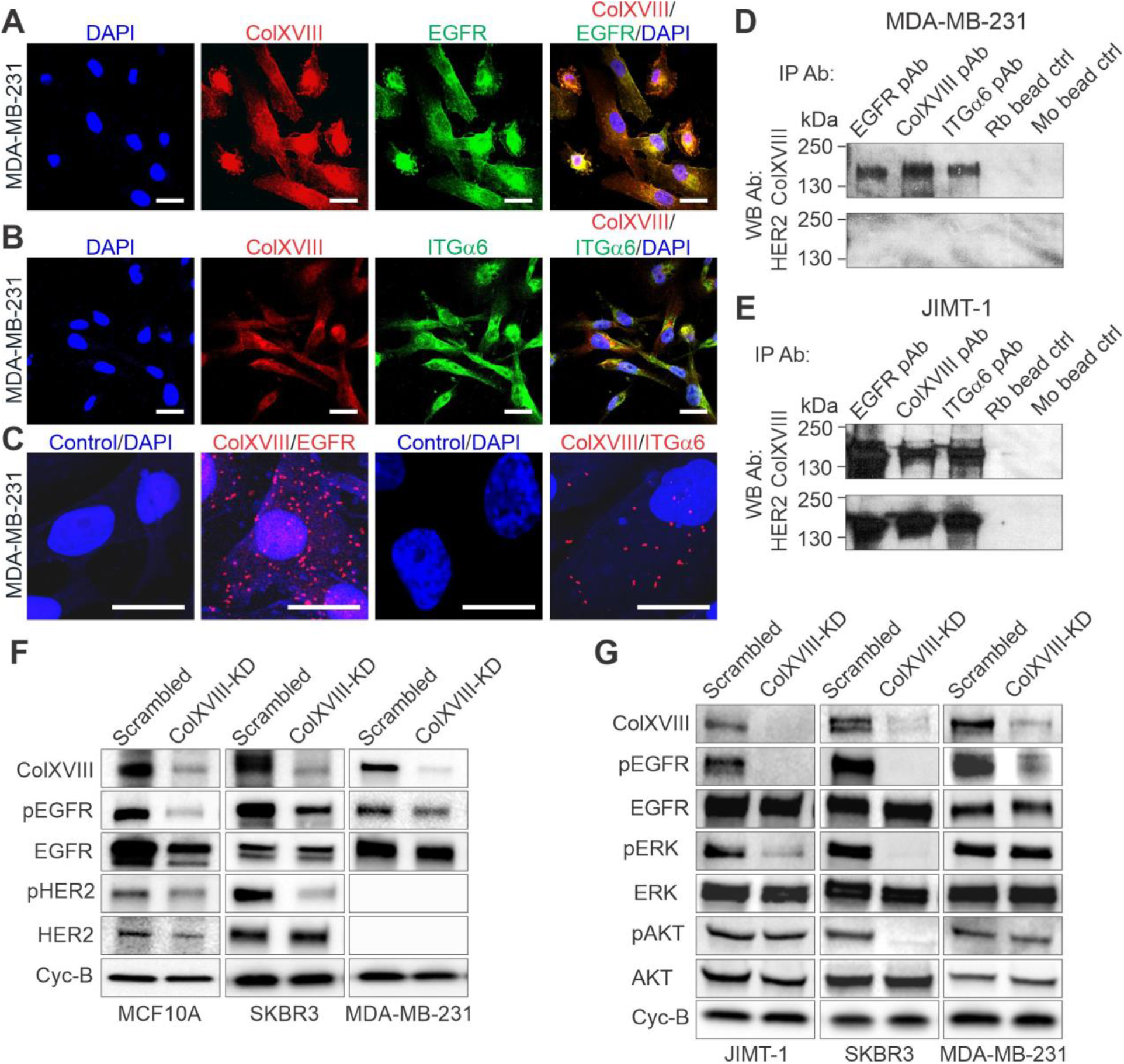
Interactions between ColXVIII, ErbBs and integrins, and analyses of EGFR/ErbB and the downstream signaling pathway. **(A-B)** Representative images of immunofluorescence staining of ColXVIII (red), EGFR (green in A) and integrin α6 (green in B) in MDA-MB-231 BC cells. Nuclei are counterstained with DAPI. Scale bars, 20μm. **(C)** *In situ* proximity ligation assay. Evidence of proximity (distance less than 40 nm) for ColXVIII (mAB DB144-N2) and EGFR (mAB 52894) (left panels), and for ColXVIII (mAB DB144-N2) and α6 integrin (pAB 97760) (right panels) is indicated by the presence of red dots. Negative controls without primary antibodies are shown. Scale bars, 20μm. **(D-E)** Co-immunoprecipitation (IP) of ColXVIII (mouse mAB DB144-N2), EGFR (rabbit mAB 52894), and integrin α6 (rabbit pAB 97760) in HER2-negative MDA-MB-231 cells and in HER2-positive JIMT-1 cells. Protein complexes were detected in Western blotting (WB) by means of ColXVIII (rabbit pAB QH48.14) and HER2 (rabbit mAB 4290) antibodies. Goat anti-rabbit (Rb) IgG and goat anti-mouse (Mo) IgG-coated magnetic bead controls are shown. **(F)** Representative immunoblots of EGFR and HER2 phosphorylation in scrambled and ColXVIII KD MCF10A, SKBR3 and MDA-MB-231 cell lysates. **(G)** Representative immunoblots of EGFR and phosphorylation of downstream signaling mediators in scrambled and ColXVIII KD JIMT-1, SKBR3 and MDA-MB-231 cell lysates.

To study further the involvement of ColXVIII in EGFR/ErbB signaling in BC and to better understand its mechanism of action at the cellular level, the effects of ColXVIII KD on EGFR/ErbBs and downstream signaling pathways, including the mitogen-activated protein kinase (Ras/Raf/MEK/ERK1/2) and phosphatidylinositol-3-kinase (PI3K/Akt) pathways, were assessed in several human BC cell lines. Western blot analyses showed that EGFR and HER2 phosphorylations were decreased in the SKBR3 and MCF10A cells upon ColXVIII KD relative to the scrambled cells, and that EGFR phosphorylation was reduced in the HER2-deficient MDA-MB-231 cell line, albeit somewhat less than in the other cell lines tested (Fig. 7F-G). Moreover, pERK and pAKT levels were decreased in SKBR3 cells (Fig. 7F-G). In the JIMT-1 cells, which have an activating mutation in the catalytic subunit alpha of the *PI3KCA* gene *(31)*, ColXVIII KD did not affect the level of pAKT, but the pERK signal was lower than in the control JIMT-1 cells. Even though pEGFR levels were reduced in the MDA-MB-231 cells, no apparent changes were observed in the phosphorylation of downstream effectors (Fig. 7F-G), probably due to activating mutations in *KRAS* and *BRAF* in this cell line *(32)*.

### Therapeutic potential of ColXVIII

In view of these results, we finally focused our interest on the potential effects of ColXVIII inhibition on drug responses when combined with the tyrosine kinase inhibitor lapatinib or with humanized mABs against HER2 (trastuzumab) and EGFR (panitumumab). Lapatinib treatment almost completely blocked the proliferation of the HER2-type SKBR3 cells, and thus ColXVIII KD, which by itself resulted in a roughly 30% reduction in cell proliferation in five days, did not yield any additional effect (Fig. 8A, Fig. 2E). SKBR3 cells responded well to HER2-targeting trastuzumab, however, and this alone led to an approximately 25% reduction in cell proliferation in five days of culture. Interestingly, simultaneous administration of ColXVIII-targeting siRNAs and trastuzumab had a synergistic effect on SKBR3 cell proliferation, leading to an over 60% reduction in cell numbers in the course of the experiment as compared with untreated scrambled cells (Fig. S9A). Moreover, EGFR-targeting panitumumab and ColXVIII siRNAs in combination inhibited the proliferation of SKBR3 cells more rapidly and efficiently than did either of these treatments alone (Fig.S9B).

**Fig. 8.**
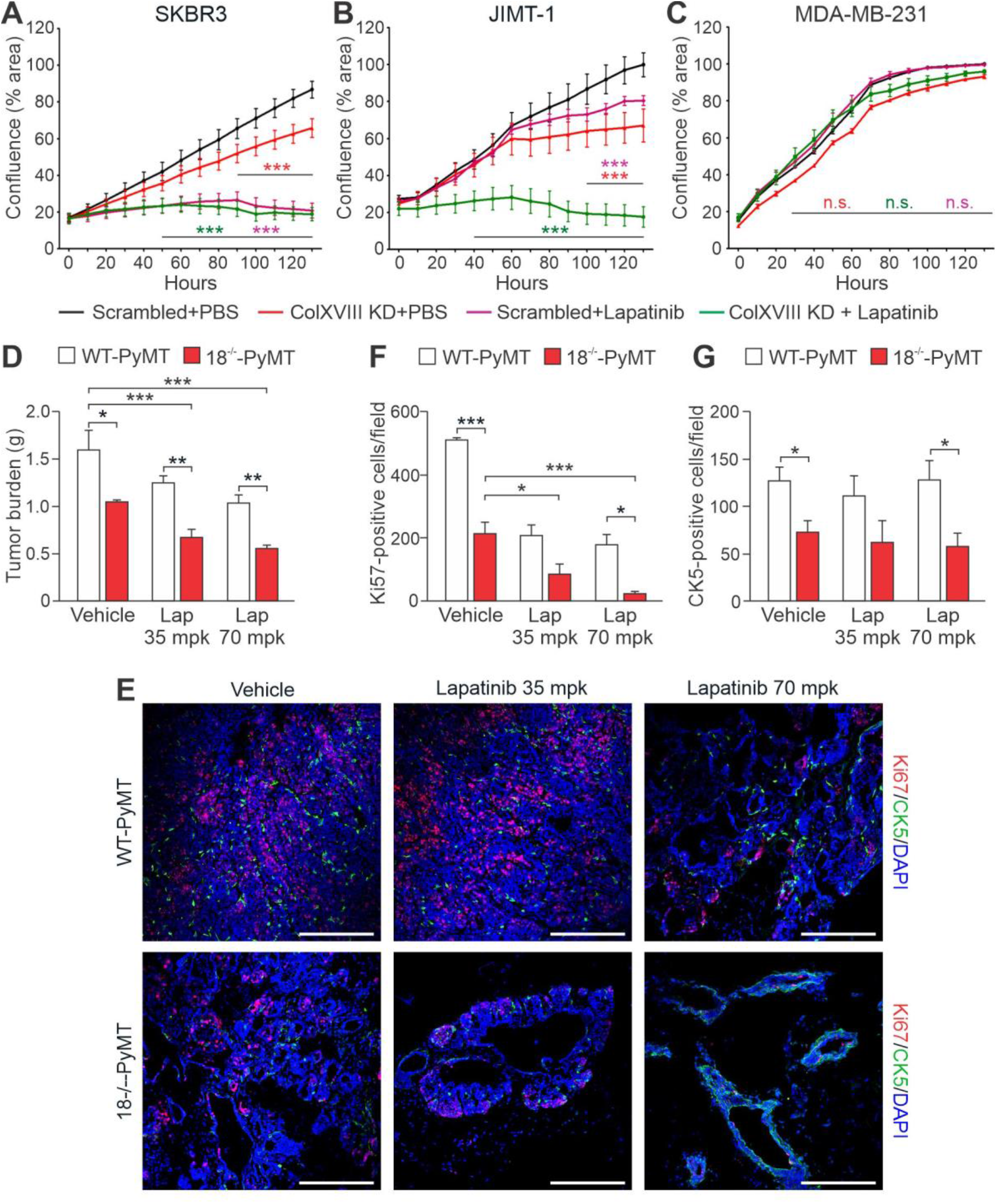
Translational potential of ColXVIII in breast cancer. **(A-C)** Ablation of ColXVIII *via* siRNA-mediated KD augments the efficacy of lapatinib, the dual inhibitor of EGFR and HER2 tyrosine kinases, in HER2-amplified SKBR3 and JIMT-1 cell lines, whereas in triple-negative MDA-MB-231 cells neither ColXVIII KD nor EGFR/HER2-targeting drugs have any effect. Cell proliferation was monitored in real time for five days using the IncuCyte live cell imaging platform, and the data are presented as percentages of confluence in the area monitored. **(D-E)** Genetic inactivation of *Col18a1* in the MMTV-PyMT mouse mammary carcinoma model augments the efficacy of lapatinib. **(D)** The total tumor burden in vehicle (0.5% hydroxypropyl methylcellulose)-treated and lapatinib-treated WT-PyMT mice and 18^−/−^-PyMT mice at the age of 10 weeks. Two doses of lapatinib, 35 mpk (milligrams per kilogram) and 70 mpk, were tested. N=6 mice per experimental group. **(E)** Representative images of proliferating Ki67-positive cells (red) and CK5-positive mammary progenitor cells (green) in the vehicle- and lapatinib-treated WT-PyMT and 18^−/−^-PyMT tumors at week 10. Scale bars, 200 nm. **(F)** Quantification of the Ki67-positive cell counts per microscopic field at 20x magnification. **(G)** Quantification of the CK5-positive cells per microscopic field at 20x magnification. N=6 mice for the Ki67 and CK5 counts, and four random fields per tumor were analyzed. In A-D and F-G, the Student’s ‘t’ test was used for the statistical analysis. *, p<0.05. **, p<0.01; ***, p<0.001; and n.s., not significant. Error bars represent s.e.m.

HER2-amplifed JIMT-1 cells are resistant to drugs that directly target ErbB receptors, due to several co-existing drug resistance mechanisms, including mutations in *PI3KCA* that activate the PI3K/AKT pathway *(31)*. We noticed that whereas neither lapatinib nor ColXVIII KD alone affected the proliferation of this cell line in the early growth phase but led to growth inhibition in the later stages, their combined effect was extremely rapid and efficient and almost completely abolished the proliferation of JIMT-1 cells (Fig. 8B). Lapatinib, panitumumab and trastuzumab treatments did not affect the proliferation of MDA-MB-231 cells because this cell line is HER2-negative and has mutations in the downstream effectors *KRAS* and *BRAF* that keep the cells in a proliferative state *(32)* (Fig. 8C, Fig. S9C-D). The EGFR-targeting panitumumab, however, did result in a significant growth restriction in MDA-MB-231 KD cells, although the effect of ColXVIII inhibition was less impressive in this cell line than in the SKBR3 and JIMT-1 cells (Fig. S9C-D). Besides these three cell lines, the HER2-positive luminal B-type BT474 cell line that has a *PIK3CA* mutation *(33)* and is thus resistant to ErbB-targeting drugs was also included in our tests. The proliferation of BT474 cells was not affected at all by trastuzumab, and only marginally by lapatinib. Depletion of ColXVIII KD alone reduced the proliferation of BT474 cells by 25-30% in five days and sensitized these cells to lapatinib (Fig. S9E-F).

Besides reducing cancer cell proliferation, ColXVIII KD slowed down the migration of SKBR3 cells significantly but, as in the proliferation assay, it did not exhibit any additional inhibitory effect on wound closure when combined with lapatinib (Fig. S9G). In MDA-MB-231 cells the inhibitory effect of ColXVIII KD was more evident in cell migration than in cell proliferation (Fig. S9H, Fig. 8C), whereas lapatinib produced only a marginal effect, as was expected due to mutations in signal mediators (Fig. S9H). The combined use of lapatinib and ColXVIII siRNAs, but not single treatments with these reagents, resulted in a significant reduction of JIMT-1 cell migration (Fig. S9I).

A preclinical *in vivo* experiment with lapatinib confirmed that ColXVIII knockout adds a significant inhibitory effect upon mammary carcinogenesis in the MMTV-PyMT mouse model. In the vehicle-treated 18^−/−^-PyMT mice the total tumor burden was approximately 30% smaller than in the vehicle-treated WT-PyMT mice (Fig. 8D). The tumor burden was further reduced in the 18^− /−^-PyMT mice treated with lapatinib, by approximately 35% and 54% compared with the vehicle-treated 18^−/−^-PyMT group, depending on the dose. Immunostainings showed that while the mammary tumors of vehicle-treated WT-PyMT mice had high numbers of proliferative Ki67-positive tumor cells, the numbers of dividing tumor cells were significantly lower in the vehicle-treated 18^−/−^-PyMT mice and in the lapatinib-treated WT-PyMT mice, and particularly low in the lapatinib-treated 18^−/−^-PyMT mice (Fig. 8E-F). Correspondingly, the tumors in the lapatinib-treated 18^−/−^-PyMT mice were considerably smaller, and those in some MGs of the 18^−/−^-PyMT mice receiving a high dose of lapatinib had been almost completely eradicated, so that the fat pads contained fairly normal-looking ductal structures (Fig. 8E). The number of intratumoral CK5^+^ progenitor cells in the 18^−/−^-PyMT tumors was initially significantly smaller than in the WT-PyMT tumors (Fig. 8E,G; Fig. 5C) and lapatinib treatment did not affect these cell counts in either genotype in the current model (Fig. 8G). In summary, our preclinical experiments demonstrate the importance of ColXVIII for BC cells functions and show that the inhibition of its action in tumor cells has important therapeutic potential.

## DISCUSSION

This study shows that ColXVIII expression is high in human and mouse BC and supports tumor cell proliferation in an autocrine manner through a previously unreported mechanism involving EGFR/ErbB signaling. Moreover, it presents evidence that ColXVIII can have significant translational value as a novel biomarker and a potential therapeutic target in BC. More specifically, our key findings are that 1) ColXVIII expression is induced in human BC cells; 2) high ColXVIII expression is associated with high-grade tumors and reduced survival, especially in the HER2 and basal/TNBC subtypes; 3) ColXVIII is co-expressed with EGFR, HER2 and integrin α6, and interacts with these receptors to induce cell proliferation and promote tumor growth, 4) ColXVIII supports stemness properties in BC cells, most likely through interactions with integrin α6; 5) the short ColXVIII isoform in particular is induced in mammary tumors; 6) the N-terminal TSP-1 domain of ColXVIII is implicated in promoting cancer cell proliferation and tumor growth; 7) ablation of ColXVIII improves the efficacy of EGFR/ErbB-targeting therapeutics in preclinical tests; and 8) plasma levels of N-terminal ColXVIII fragments are higher in BC patients than in healthy controls.

Our *in vivo* studies using the MMTV-PyMT mammary cancer model crossed with our unique total and isoform-specific *Col18a1* knockout models convincingly demonstrated for the first time that the short ColXVIII is the key isoform upregulated in mammary tumors. Importantly, the short isoform was found to be responsible for the pro-tumorigenic action of ColXVIII, as the specific deletion of this isoform, but not the medium/long isoforms, significantly inhibited cancer cell proliferation, the primary tumor burden and lung metastasis (Fig. 3, Fig. S6). The short isoform has a TSP-1 domain at the N-terminus of the molecule, and it is shown here that recombinant fragments containing the TSP-1 sequence can at least partially recover the proliferation deficit caused by ColXVIII KD in human BC cells, whereas C-terminal endostatin proved ineffective in mediating this task (Fig. 2E-G). The current understanding of the specific functions of ColXVIII isoforms and their N-terminal NC domains is still limited and is focused on their developmental functions [as summarized in *(20)*]. Thus our finding that the short ColXVIII and its TSP-1 domain have pro-tumorigenic functions is a pioneering discovery.

This is also the first study to demonstrate that ColXVIII is co-expressed and forms a complex with ErbB receptors and integrin α6 in BC cells, thus having the potential to enable downstream signaling through MAPK/ERK and PI3K/AKT pathways, leading to increased tumor cell proliferation and migration. The concept of integrated signaling through growth factor and ECM receptors is well established, and the downstream pathways of the two signaling systems overlap inside the cells *(34)*. At present, however, we have no experimental data to prove whether ColXVIII binds directly to the EGFR/HER2 receptor pair and/or the α6 integrin, but we speculate that ColXVIII might be involved in coordinating the growth factor and ECM receptor signaling events.

We can compare our data on ColXVIII with findings regarding other ECM molecules highly expressed in cancer cells and implicated in EGFR/ErbB signaling. Tenascin-C, for example, resembles ColXVIII in many ways: it is commonly upregulated in cancer cells, especially at the invasive tumor front, it associates with a poor clinical outcome, and binds integrins and EGFR through EGF-like repeats to induce tumor cell proliferation and invasion and to support CSCs *(35– 38)*. Laminin 332 is an example of a BM protein interacting with both EGFR and integrins to sustain tumorigenesis. ECM remodeling in cancer reveals cryptic EGF-like repeats from laminin 332 which is thought to stimulate SCC tumorigenesis in an EGFR- and α6β4 integrin-dependent mechanism *(38–40)*. ECM proteins lacking the EGF repeats, such as decorin, can also bind to and activate EGFR and other ErbBs, but unlike tenascin-C and laminin 332, decorin displays anti-tumor activities *(41, 42)*. Our novel data thus place ColXVIII on the list of ECM components with a role in modulating EGFR/ErbB signaling in cancer.

Our analyses of public databases (Fig. 1, Fig. S2, Fig. S3) and human BC tissue and liquid biopsy samples (Fig. 6, Fig. S4) showed that high ColXVIII expression is associated with a bad prognosis, especially in the aggressive HER2 and basal/TNBC subtypes. These observations sustain the hypothesis that ColXVIII is implicated in BC progression and that its targeting could be beneficial in improving treatment outcomes in combination with drugs targeting these GFRs. This assumption was confirmed by experiments showing that ColXVIII deprivation in HER2-positive BC cells can enhance the efficacy of EGFR/ErbB-targeting drugs, both *in vitro* and *in vivo* (Fig. 8, Fig. S9). Simultaneous ColXVIII and EGFR/ErbB targeting was furthermore shown to be beneficial in the lapatinib-resistant HER2-positive BT474 cell line, which has an activating *PIK3CA* mutation downstream of the EGFR/HER2 receptor complex *(33)* (Fig. S9F), indicating that ColXVIII inhibition can alleviate drug resistance.

Biochemical and biomechanical signals from the three-dimensional ECM are implicated in the response and resistance of cancer drugs *(30, 43, 44)*. Mechanisms by which the inhibition of ColXVIII can overcome resistance to EGFR/ErbB-targeting drugs in HER2 type BC cells (Fig. 8, Fig. S9) can be speculated upon in the light of our observations and data concerning other ECM molecules. It has been reported that disruption of the interaction between laminin 332 and integrins α6β4 or α3β1, and thereby cell adhesion to the BM, can sensitize HER2-positive BC cells to trastuzumab and lapatinib treatments by inhibiting the PI3K/AKT, MAPK/ERK1/2 and focal adhesion kinase (FAK) pathways *(45)*. In other studies, high β1 integrin expression has been shown to predict a poor prognosis for trastuzumab- and lapatinib-treated HER2-positive BC and induce resistance to these drugs through FAK and Src signaling *(46, 47)*. The same treatments have been shown to induce expression of several ECM genes through β1 integrin and Src, including the aforementioned decorin and tenascin-C as well as many collagens *(48)*. Interestingly, ECM stiffness per se reduces drug and radiation sensitivity in many cancers, e.g. by forming a physical barrier against drug infiltration and by CSC promotion via various molecular mechanisms, including the regulation of integrin signaling *(43, 49, 50)*.

As a ubiquitous niche component, ColXVIII has roles in maintaining various types of tissue stem and progenitor cells [as summarized in *(22)*]. We have shown previously that the N-terminal sequences in the medium/long ColXVIII isoforms in adipose tissue support the differentiation of progenitor cells/committed precursors to form mature adipocytes *(51)*. Studies by Gupta *et al*. revealed that ColXVIII is overexpressed in therapy-resistant breast CSCs, suggesting that it may have a role in the generation and propagation of these cells *(25)*. Our work provides additional experimental evidence to support this finding, since the numbers of cells with CSC characteristics were reduced both in mouse mammary tumors with *Col18a1* deletion and in human BC cells with reduced ColXVIII expression (Fig. 5, Fig. S7). The demonstrated interaction between ColXVIII and integrin α6 in BC cells (Fig. 7), a key integrin subunit in breast CSCs *(26)*, is probably implicated in the maintenance of the stemness properties of BC cells. CSCs are not only responsible for sustaining primary tumors, but are also connected with the metastatic dissemination of neoplastic clones to distant organs *(12–15)*, and we show that both the primary tumor burden and lung colonization are reduced in mice with full or partial depletion of *Col18a1* isoforms (Fig. 3, Fig. S6). Our previous work has shown that deletion of *Col18a1* leads to BM loosening and reduced stiffness *(52)*. Thus, it is possible that ColXVIII upregulation in solid cancers may affect both the biomechanical properties of the tumor ECM and the maintenance of CSCs, and thereby regulates tumor promotion and drug responses.

In conclusion, our findings indicate that signaling cues triggered by the N-terminal TSP-1 domain of ColXVIII and transmitted through EGFR/ErbBs and/or integrins can potentiate BC cell functions and promote the development of drug resistance, especially in the advanced HER2-type BC. The targeting of ColXVIII in the TME could therefore provide a novel therapeutic approach for achieving BC regression, even in cases where the tumor does not show any response to the clinically tested drugs that inhibit EGFR/ErbB signaling. Our data also show that ColXVIII could be of substantial value as a biomarker of BC progression, either scored in tissues, or observed in liquid biopsies even before metastasis.

## MATERIALS AND METHODS

### Study design

The objective of this study was to examine the expression, roles, mechanisms of action and prognostic and therapeutic relevance of the BM component ColXVIII in BC.

### Human BC samples and survival analysis

The expression and localization of ColXVIII in 116 human BC tissue samples was analyzed by IHC and circulating levels of ColXVIII were determined in 32 BC patients and six healthy volunteers by ELISA assay. All the human samples and clinical data were anonymized and labelled with a research code for blinded histopathological and plasma analyses. Only authorized personnel of the Oulu and Umeå University Hospitals had access to personal and clinical data. Informed consent for data use was obtained from all the patients. The Ethical Committee of the Northern Ostrobothnia Health Care District (Dnr 88/2000 and amendment, Dnr 194/2013, Dnr 100/2016) and the Ethical Committee at the Medical Faculty of Umeå University (Dnr 09-175M) granted ethical approval for the use of these samples. The association of ColXVIII expression with BC patient survival was analyzed in open databases using the Kaplan Meier survival analysis at the website www.kmplot.com *(24)*.

### Mouse models

To study the functions of ColXVIII in BC *in vivo*, a transgenic mouse mammary carcinogenesis model based on mammary tumor virus promoter-driven expression of the polyoma middle T antigen (MMTV-PyMT) was crossed with three different *Col18a1* knockout models. All the animal experiments were approved by the Finnish National Animal Experiment Board (permits ESAVI/6105/04.10.07/2015, ESAVI/1188/04.10.07/2016 and ESAVI-2936-04.10.07/2016) and conducted at the University of Oulu Laboratory Animal Centre. Mammary tumor growth was monitored in randomized experimental groups of females at specific time intervals depending on the rate of tumor growth and the humane endpoint criterion as explained in the Supplement. The number of animals per genotype at various time points ranged from a minimum of three mice at week 6 up to 14 mice at week 13. Lung metastasis was studied in experimental groups aged 10−12 weeks, 13−15 weeks and 16−18 weeks, ten mice in each group per genotype. To study the tumor-cell autonomous vs. microenvironmental role of ColXVIII, a reciprocal orthotopic allograft transplantation experiment between control and *Col18a1*-deficient mice (N=6−12) was performed. *In vivo* animal experiments were conducted in a non-blinded manner. Mouse tumor tissues were studied by histological and immunohistochemical, morphometric, flow cytometric and qRT-PCR methods. The numbers of samples and replicates for the quantitative analyses are indicated in the Figure legends.

### Cell studies

Expression of ColXVIII in the human BC cell lines was analyzed with qRT-PCR, Western blotting and immunocytochemistry. *In vitro* loss-of-function experiments were performed by siRNA-based KD of ColXVIII in BC cells, and gain-of-function experiments were conducted with ColXVIII KD cells supplemented with recombinant ColXVIII fragments. Live cell imaging was used for functional analyses of cell proliferation and migration. In order to study ColXVIII-related signaling, the interactions of ColXVIII with EGFR/ErbB and integrin receptors were examined using proximity ligation and co-immunoprecipitation experiments, and phosphorylation of EGFR/Erbs and the downstream signaling mediators was assessed by Western blotting. Cell culture experiments were performed independently at least twice per experiment as indicated in the Figure legends.

### Drug tests

To investigate the medical relevance of ColXVIII targeting in combination with current BC therapies, the EGFR/ErbB inhibitors lapatinib, trastuzumab and panitimumab were used in combination with ColXVIII KD in human BC cell lines cells and their respective controls. *In vivo* lapatinib treatment was applied to six control and six *Col18a1* null PyMT mice using two different doses to test the efficacy of the drug.

### Statistical Analysis

Statistical analyses were performed using the unpaired ‘t’ test for experiments with two groups, and a two-way analysis of variance test (Bonferroni’s posttest) when comparing data from experiments with multiple groups. A repeated measures one-way analysis of variance was used to analyze the primary tumor growth curves (Dunnett’s multiple comparison test and Bartlett’s post-correction test). Mouse survival analysis was performed using the log rank (Mantel–Cox) test. Differences were considered statistically significant at a p-value less than 0.05. GraphPad Prism software was used for the statistical analyses.

Further details of the materials and detailed experimental protocols are presented in the Supplementary Materials and Methods.

## Acknowledgments

We thank Jaana Peters, Annette Berglund, Päivi Tuomaala, Sirkka Vilmi and Aila White for their excellent technical assistance; Dr. Virpi Glumoff for support in FACS analyses; Biocenter Oulu Tissue Imaging Center and Biocenter Finland, especially Dr. Veli-Pekka Ronkainen and Antti Viklund, for support in microscopy; and Oulu Laboratory Animal Centre and Biocenter Transgenic and Tissue Phenotyping Core Facility for the facilities, services and assistance in mouse work.

## Author contributions

R.D. designed and performed all the experiments, analyzed the data, and wrote the manuscript.

H.P. supervised the work, analyzed the human IHC data and wrote the manuscript.

V.I. designed some of the experiments, analyzed the data and provided scientific input

G.R., M.R.V. and T.V. scored the human IHC.

H.R. designed and performed the mouse experiments.

S.K. provided the human BC samples and analyzed the human IHC data.

G.M.N. performed some of the mouse experiments.

S.M.K. analyzed ColXVIII expression in the human BC samples.

I.K. produced the recombinant proteins.

J.K. provided scientific input.

T.S. provided the mouse ColXVIII antibody.

R.W. provided human tissue samples.

F.W. provided human samples and scientific input.

M.S. provided human samples and scientific input, and wrote the manuscript.

T.P. conceptualized and supervised the studies, provide scientific input, and wrote the manuscript.

R.H. conceptualized and supervised the studies, provided scientific input, and wrote the manuscript.

## Data and materials availability

The data associated with this study are present in the paper or the Supplementary Materials. Materials are available upon request from the corresponding author.

