## Supplementary material for "Collagen XVIII promotes breast cancer through EGFR/ErbB signaling and its ablation improves the efficacy of ErbB-targeting inhibitors"

**Devarajan et al.**

**Supplementary Materials:**

**Materials and Methods**

**Supplementary Figures**

Fig. S1. ColXVIII immunohistochemistry in human breast cancer and the normal mammary gland

Fig. S2. Association between ColXVIII expression and patient survival

Fig. S3. ColXVIII expression and patient survival rates in grade 3 breast tumors of different molecular subtypes

Fig. S4. ColXVIII in breast cancer patient plasma

Fig. S5. The short ColXVIII isoform supports mammary carcinogenesis in the MMTV-PyMT model

Fig. S6. Lung metastasis in wild type MMTV-PyMT and ColXVIII-deficient PyMT mice

Fig. S7. ColXVIII supports breast cancer stem cells

Fig. S8. Interactions between ColXVIII, ErbBs and integrins

Fig. S9. Analysis of breast cancer cell proliferation and migration upon ColXVIII knockdown and/or EGFR/ErbB-targeting drug treatments

**Supplementary Tables**

Table S1. Human breast cancer samples with histopathological and relapse data

Table S2. Antibodies

Table S3. qRT-PCR primers

### Materials and Methods

#### Human breast cancer (BC) samples and ethical permits

Formalin-fixed paraffin-embedded (FFPE) human BC samples representing different subtypes were received from two patient cohorts, at Oulu University Hospital, Finland (n=21), and Umeå University Hospital, Sweden (n=95) (Table S1). The plasma samples from BC patients (n=8-12 per molecular subtype) were from the Umeå University Hospital cohort. The control plasma samples were collected from 6 healthy women volunteers above 45 years of age. All human samples and clinical data were anonymized and labelled with a research code for blinded histopathological analyses. Only authorized personnel of the Oulu and Umeå University Hospitals had access to the personal and clinical data. Informed consent for the data was obtained from all the patients. The Ethical Committee of the Northern Ostrobothnia Health Care District (Dnr 88/2000 and amendment, Dnr 194/2013, Dnr 100/2016) and the Ethical Committee at the Medical Faculty of Umeå University (Dnr 09-175M) granted approvals for the use of these samples.

#### Immunohistochemistry of human breast cancer samples

Immunohistochemical (IHC) staining of 5 μm thick FFPE breast tumor sections was performed using the Dako REAL™ EnVision™ detection system (Dako, #K5007). The primary antibodies (AB) used were an in-house mouse monoclonal antibody (mAB) DB144-N2 which targets the N-terminal portion of the molecule of human ColXVIII *(53)*, dilution 1:150, an in-house rabbit polyclonal antibody (pAB) QH48.18 which targets the same N-terminal region of human ColXVIII *(54)*, dilution 1:000, rabbit mAB for human EGFR (Cell Signaling Technology, #4267), dilution 1:50, rabbit mAB for human HER2 (Cell Signaling Technology, #4290), dilution 1:50, and mouse mAB for human Ki67 (Dako, #M7248), dilution 1:50 (Table S2). Images were captured using a Leica DM LB2 light microscope or automatic scanning using a Zeiss Axio Imager.M2m. The ColXVIII, EGFR, HER2 and Ki67 expression analyses of the IHC stainings were performed by two independent observers, and the semi-quantitative scores for ColXVIII and EGFR are reported as averages of the cytoplasmic staining intensity in tumor cells (0=negative, 0,5-1=weak, 1,5-2=moderate, and 2,5-3=strong). The consistency of the scoring was tested using linear regression or Pearson correlation, and non-matching scores were removed from the final analysis.

**Kaplan-Meir survival analysis**

Kaplan Meier survival analysis was performed by accessing BC patients' data for *COL18A1* gene expression (Affymetrix *COL18A1* probe: 209082_s_at 'TCAGCCAGCACTTGTCCAGCTGAGC' targeting the endostatin domain) using the website *(24)*. This database provides a compilation of data from several open databases, including the Cancer Genome Atlas (TCGA), the Gene Expression Omnibus (GEO and the European Genome-phenome Archive (EGA). The association of ColXVIII expression levels with overall survival (OS), relapse free survival (RFS) and distant metastasis-free survival (DMFS) was analysed either in unclassified form or classified according the differentiation grade or molecular subtype. The auto-select best cutoff was used for calculating the survival plots, and false discovery rate was less than 5% in all analyses.

**Enzyme-linked immunosorbent assay (ELISA) for plasma samples**

Indirect ELISA was carried out to detect the N-terminal ColXVIII fragments in blood plasma samples collected from BC patients and healthy controls. 100µl of plasma samples, diluted 1:10 in 100 mM carbonate/bicarbonate coating buffer, pH 9.6, were pipetted in duplicate onto a 96-well ELISA microplate (Nalgene Nunc, USA). Linear dilutions (1-300 ng/ml in coating buffer) of the standard, the recombinant N-terminal NC11 fragment of ColXVIII *(51)*, were added in duplicate. The plates were incubated overnight at 4°C with gentle shaking and washed thrice with PBS-0.05% Tween (PBST). The remaining protein binding sites in the coated wells were blocked by adding 200µl of blocking buffer (1% BSA in PBST) and incubating at +37°C for 1h. After washing the wells thrice with PBST, 100µl of monoclonal DB144-N2 *(53)* antibody against the TSP-1-like domain of ColXVIII[1], diluted 1:1000 in blocking buffer, was added and incubation was continued for 1 h at +37 °C. After washing thrice with PBST, an anti-mouse HRP-conjugated secondary antibody at a dilution of 1:10 000 was added to each well and incubated for 1 hr at +37°C. Control wells without primary or secondary antibodies were included in the assay. After washing thrice with PBST, 100µl of freshly constituted 3,3',5,5'-tetramethylbenzidine (TMB) substrate was added and the reaction was allowed to proceed at room temperature without shaking. The colour reaction was stopped after 15-20 min by adding an equal volume of 1M $H_2SO_4$. The absorbance was read at a wavelength of 450 nm and a correction wavelength of 650 nm using a Tecan Infinite M1000 PRO microplate reader. The concentration of ColXVIII in the plasma samples was determined from the standard curve and the results were analysed using Graph-pad Prism statistical software and represented in graphs.

**Human breast cancer cell lines**

MDA-MB-231 (HTB-26) and MCF7 (HTB-22) cells were purchased from the American Type Culture collection, MCF10A (CLL1040) cells from Sigma-Aldrich, and JIMT-1 (ACC589) cells from DSMZ, Germany. SKBR3, Hs578T and T47D cells were a kind gift from Professor Peppi Koivunen, University of Oulu, and BT474 cells from Dr. Jussi Koivunen, Oulu University Hospital. Cell lines other than MCF10A were cultured in Dulbecco's Modified Eagle's Medium (DMEM) with high glucose and supplemented with 10% fetal bovine serum (FBS), 1% penicillin/streptomycin and 1% GlutaMax (all from Gibco). MCF10A cells were cultured in DMEM:F12 medium containing 5% horse serum (Invitrogen 16050-122) FBS, 10 μg/ml of insulin (Sigma-Aldrich #I9278), 20 ng/ml of EGF (Sigma-Aldrich #E9644), 100 ng/ml of cholera toxin (Sigma-Aldrich #C8052), 0.5 μg/ml of hydrocortisone (Sigma-Aldrich #H0888) and 1% penicillin/streptomycin. All the cell lines were maintained at 37$^{o}$C with 5% ambient $CO_{2.}$ and found negative in routine mycoplasma infection tests.

**siRNA-mediated *COL18A1* gene knockdown**

Transfections were performed by first incubating 500µl of Opti-MEM media (Gibco) with 7.5µl of lipofectamine (RNAiMAX, Invitrogen) in each well of the 6-well plate for 5 min at RT. After that 10nM of siRNAs were added, mixed gently and incubated for an additional 15 min at RT. BC cells were added to the transfection mixture at a density of $2.5 \times 10^5$ cells per well. For an efficient knockdown of the ColXVIII, a mixture of two commercially available siRNAs targeting two distinct areas in the ColXVIII mRNA transcript were used, namely SASI_Hs01_00161793 (Sigma-Aldrich) for the 5′- GCATCTTCTCCTTTGACGGCAAGGA-3′ sequence in the N-terminal TSP-1 domain, and Hs_COL18A1_3 (Qiagen) for the sequence 5′-

CCCGGGATGAACGGATTGAAA-3′ in the C-terminal endostatin domain. 10nM of scrambled siRNA with a random nucleotide sequence (AllStars Negative Control siRNA, Qiagen) was transfected into cells in the same way and used as a negative control. Knockdown efficiency was estimated with qRT-PCR for the ColXVIII mRNA transcripts and Western blot for the ColXVIII protein. Transfection efficiency was ensured with Allstar Hs Cell Death Control siRNA (Qiagen), which kills the cells that have become transfected. The transfected cells were further seeded at a density of $5\times10^3$ or $3–4\times10^4$ cells/well in a 96-well plate, depending on the protocol.

**Cell proliferation and migration assays**

For the cell proliferation and migration analyses, live cell phase contrast imaging was performed with IncuCyte ZOOM (Essen Biosciences). For the proliferation assays $5\times10^3$ cells/well were plated onto 96-well cell culture plates and cell proliferation curves were drawn from confluence measurements acquired during round-the-clock kinetic imaging at 2-hour intervals. To study the effects of specific non-collagenous (NC) domains of ColXVIII on cell proliferation, recombinant TSP-1 *(55)*, endostatin *(56)* and the NC11 fragment of the long ColXVIII *(51)* were produced in mammalian cells and added to cells at final concentrations of 500 ng/ml. In the cell migration assays $3\text{-}5\times10^4$ cells/well were plated onto 96-well ImageLock plates (Essen Bioscience). At 90–100% confluence the plates were scratched with a 96-well wound maker (Essen Bioscience) and cell migration was detected with IncuCyte by recording one image per well every two hours for 30-96 hours. Cell proliferation was quantified by determining the confluence of the surface area of the 2D culture over time and cell migration by monitoring wound closure using the Incucyte zoom software. Statistical analyses were performed with Graph-pad Prism software.

**Mouse models and ethical permits**

Previously generated *Col18a1*$^{-/-}$ knockout mice (*Col18a1*$^{tm1Hms}$, MGI:2179134) lacking all three ColXVIII isoforms *(57)*, *Col18a1*$^{P1}$ mice (*Col18a1*$^{tm1Pih}$, MGI:4948124) lacking the short ColXVIII isoform *(51)* and *Col18a1*$^{P2}$ mice (*Col18a1*$^{tm2Pih}$, MGI:4948125) lacking the medium/long isoforms *(51)* were used. All three ColXVIII mouse lines were backcrossed into the FVB/N (Harlan, the Netherlands) background for at least 8 generations. These mouse lines were then intercrossed with the transgenic MMTV–PyMT mouse line [FVB/N-Tg (MMTV-PyMT)634Mul/J, MGI:2161653]*(58)* (Jackson Laboratory, USA) for mammary carcinogenesis studies. All the animal experiments were approved by the Finnish National Animal Experiment Board (permissions ESAVI/6105/04.10.07/2015, ESAVI/1188/04.10.07/2016 and ESAVI-2936-04.10.07/2016) and conducted in the University of Oulu Laboratory Animal Centre.

**MMTV-PyMT mammary carcinogenesis studies**

The MMTV-PyMT mouse lines were maintained by breeding the PyMT transgene-positive male mice, either wild type (PyMT) or various *Col18a1* knockout males (i.e. $Col18a1^{-/-}$, $Col18a1^{P1/P1}$ and $Col18a1^{P2/P2}$ males) with PyMT-negative wild type FVB/N females or PyMT-negative knockout females to obtain female mice that were hemizygous for the PyMT transgene for use in the mammary carcinogenesis experiments. The resulting mouse lines are referred to in this paper as WT-PyMT (controls) and $18^{-/-}$-PyMT, P1-PyMT and P2-PyMT. The mice were group-housed in corncob bedding with enrichments, given purified water ad libitum and irradiated standard rodent chow, and maintained on a 12:12-h light: dark cycle. All the data reported here refer to female mice monitored at specific time intervals starting at the age of five weeks and continuing up to 14 weeks of age for the WT-PyMT mice and up to 18 weeks for the $18^{-/-}$-PyMT mice. The mice were sacrificed with $CO_2$ inhalation and cervical dislocation at predetermined time points, or upon reaching a humane endpoint criterion (size of a single mammary gland over 1 cm in diameter, high total tumor burden, loss of weight, reduced mobility, hunched posture, low activity, appetite loss, or closed eyes). The mammary glands were then dissected and weighed. For further analyses, the mammary glands were fixed with 4% paraformaldehyde (PFA) in phosphate-buffered saline (PBS), pH 7.4, and embedded in paraffin, or directly embedded in O.C.T. cryo-compound (Tissue-Tek OCT, Sakura) and stored at -80°C, or else snap-frozen in liquid nitrogen for protein and RNA analyses. Survival analysis for the WT-PyMT (n=38), $18^{-/-}$-PyMT (n=31), P1-PyMT (n=28) and P2-PyMT (n=33) mice was performed by following the subjects until the humane end point (code=1) or ruled out for noncompliance (code=0) and calculating the log rank (Mantel–Cox) test using Graph-pad Prism software.

**Mouse tissue stainings**

The tissue morphology of the mammary gland tumors was assessed in hematoxylin-eosin-stained 5 μm sections from the PFA-fixed tumor samples. For immunofluorescence staining, the PFA-fixed samples were deparaffinized for 3 min in xylene and then rehydrated with decreasing ethanol concentrations of 95, 85 and 70% and finally PBS, twice for 3 min in each solution. Heat-mediated antigen retrieval was accomplished by boiling the sections in 0.1M citrate buffer, pH 6.0, for 20 min. The sections were then blocked with 1% BSA + 0.3M glycine in a 1×Tris-buffered saline, 0.1% Tween-20, pH 7.5 (TBST) buffer followed by overnight incubation at +4°C with polyclonal antibodies) for mouse ColXVIII (anti-NC11 *(59)*, Ki67 (Dako, #7249), cleaved caspase 3 (R&D systems, #MAB835), cytokeratin 5 (CK5, Covance #PRB-160P), integrin α6 (Abcam, #181551) and integrin β1(Millipore #MAB1997) (Table S2). After incubation with the primary antibodies, the slides were washed thrice for 5 min in PBS, followed by incubation with respective fluorochrome-conjugated Alexa Fluor 488- or Cy3-conjugated secondary antibodies and DAPI (Sigma-Aldrich) for 1 hour at RT. After three 5-min washes with PBS, the samples were mounted in Immu-Mount medium (Thermo Scientific). In the case of Cy3-conjugated αSMA (Sigma-Aldrich, #C6198) the secondary antibody was omitted. 7-μm cryosections on poly-L-lysine coated slides were fixed for 20 min in methanol at –20°C, air-dried, washed for 5 min in PBS and then processed according to the above protocols. Images were acquired using Zeiss LSM 780 and LSM 700 confocal microscopes, and image analyses were performed using Fiji ImageJ or Zen 2011 software. Morphometric analyses for CK5 and Ki67 stainings were performed by capturing the images with a 20× objective using a Zeiss LSM 700/780 confocal microscope and counting positive cells with the Fiji ImageJ software. The count of positive cells was reported as the average of 4 areas of the tumor tissue.

**Carmine Alum staining of mouse mammary glands**

Carmine Alum staining was performed as previously described *(60)*. Briefly, whole mammary glands were mounted on a microscope slide, fixed in 4% PFA in PBS, pH 7.4, overnight, and transferred to 70% ethanol. The mounted slides were then rinsed in water and stained in a filtered solution of 0.2% Carmine (Sigma-Aldrich) and 0.5% aluminum potassium sulfate dodecahydrate

(Sigma-Aldrich) as previously described[6]. The samples were dehydrated through sequential 70, 95 and 100% ethanol washes (20 min in each) and stored in a methyl salicylate solution.

**Orthotopic allograft transplantation assay**

The allograft transplantation assay was performed as a reciprocal transplantation of WT-PyMT and $18^{-/-}$-PyMT tumor cells into mammary fat pads of WT and *Col18a1*$^{-/-}$ FVB mice at 5 weeks of age. The WT-PyMT and $18^{-/-}$-PyMT donor mice were euthanized and sterilized with 70% ethanol. The mammary glands were exercised and placed onto a tissue-culture plate containing DMEM:F12 medium (ATCC cat no. 30-2020) and extra fat was carefully removed and the tumors minced with scalpel blades inside laminar hoods. The minced tissues were transferred to 0.3% collagenase (LS004176, Worthington) and incubated for one hour at +37°C. The cell suspension was filtered through a 70 μm cell strainer (BD Biosciences), centrifuged and cell pellets isolated, suspended and passed again through a 70 μm cell strainer. The tumor cell suspension was reconstituted at $5×10^6$ cells/ml in PBS and 50µl of this suspension ($2.5×10^5$ tumor cells) was injected into abdominal mammary fat pads. The experimental groups were the following: Group-1, WT- FVB mice receiving WT-PyMT tumor cells; Group-2, WT-FVB mice receiving $18^{-/-}$-PyMT tumor cells; Group-3, *Col18a1*$^{-/-}$ FVB mice receivinged WT-PyMT tumor cells; and Group-4, *Col18a1*$^{-/-}$ FVB mice receiving $18^{-/-}$-PyMT tumor cells. The tumors were measured with a caliper twice per week, and their volumes calculated using the formula: length × width$^2$ × 0.52. The experiment was terminated at week 10, when the WT-PyMT tumors reached an average volume of 600 mm$^3$, or at week 17 in the case of the $18^{-/-}$-PyMT tumors, when the largest tumors reached an average of 400 mm$^3$.

**RNA isolation and qPCR**

RNA from human cell lines and mouse tissue samples was isolated using the Qiagen kit (RNeasy mini), and cDNA was produced with the iScript cDNA synthesis kit (Bio-Rad). Quantitative real-time PCR (RT-qPCR) was performed with SYBR Green Supermix (Bio-Rad). All the assays were performed in duplicate using a CFX96 Real-Time System (Bio-Rad). The values for all the samples were normalized to *Gapdh* (mouse) or *GAPDH* (human), and the relative fold change

($2^{\Delta\Delta}$Cq) was calculated using CFX Manager Software (Bio-Rad). The qRT-PCR primers used here are listed in Table S3.

**Western Blotting**

To prepare the human BC cell lysates, the culture medium was aspirated, and cells were washed once in ice-cold PBS, after which NETN-300 buffer (20 mM Tris-HCl, pH 7.5, 300 mM NaCl, 1 mM EDTA, 0.5% NP-40) supplemented with 1× complete mini EDTA protease inhibitor (Roche Diagnostics) was pipetted directly onto the plate before collecting the cells using a cell scraper. Cell lines and tumor tissue lysates were cleared by centrifugation at 20,000 × g for 20 min at 4°C and the protein concentration was measured using a DirectDetect IR spectrometer (Merck Millipore). The lysates were suspended in 2.5× Laemmli buffer, separated using SDS-PAGE and blotted onto PVDF membranes. The membranes were then washed in 1×Tris-buffered saline, 0.1% Tween-20, pH 7.5 (TBST) and blocked with a buffer containing 1% BSA and 1% skimmed milk in TBST). This was followed by incubation with primary antibodies (Table S2) overnight at +4 $^{o}$ C, three washes with TBST and incubation with horseradish peroxidase-conjugated secondary antibodies (Jackson Immunoresearch) for 2 hours at RT. Signals were detected using SuperSignal™ West Pico Chemiluminescent Substrate (ThermoFisher Scientific) following the manufacturer's instructions and detecting the chemiluminescence with a Fujifilm LAS-3000 imaging system.

**Proximity Ligation Assay**

The proximity ligation assay (PLA) to study potential interactions between ColXVIII, EGFR and integrin $\alpha$6 in cultured MDA-MB-231 and JIMT-1 cells was performed with the Duolink Detection Kit (#DUO92101, Sigma-Aldrich). The cells were seeded at a density of $1\times10^4$ per well in an 8-well Nunc Lab-Tek II Chamber Slide System (ThermoFisher) and grown overnight. They were then briefly washed with PBS, followed by fixation in ice-cold 4% PFA for 10 min and permeabilized with 0.1% triton X-100 in PBS for 10 min. The cells were washed thrice for 5 min with PBS, after which the PLA protocol described by the manufacturer was followed (Duolink® PLA Fluorescence Protocol, SigmaAldrich). The primary antibodies used in the PLA assay were mouse monoclonal AB (mAB) DB144-N2 *(53)* for ColXVIII, dilution 1:150, rabbit mAB for

EGFR (Abcam #52894), dilution 1:100, and rabbit polyclonal AB (pAB) for α6 integrin (Abcam, #97760), dilution 1:100 (Table S2). The slides were mounted in Duolink® PLA Mounting Medium with DAPI, and the images were captured using a Zeiss LSM700 Microscope with a 100× oil immersion objective.

**Co-immunoprecipitation**

Whole-cell lysates prepared in NETN-300 buffer from a sub-confluent 10 cm dish of MCF10A, SKBR3, JIMT1 and MDA-MB-231 cells were used for immunoprecipitation with antibodies against ColXVIII, EGFR and HER2, and integrins α6 and β1. For immune complex formation with the target molecule in the lysate, 5 μg of anti-EGFR (Abcam #52894), anti-ColXVIII (DB144-N2 or QH48.14) *(53, 54)*, anti-α6 integrin AB (Abcam, #ab97760), or anti-β1-Integrin (Santa Cruz SC9970) antibody (Table S2) was incubated with 1 ml of cell lysate overnight at +4°C with gentle rocking. The immune complex was then captured on Protein A- or G-coupled magnetic beads (Dynabeads, Invitrogen) and incubation was continued overnight at +4°C. The protein complex bound on the beads was pelleted with a magnetic separation rack and the beads were washed thrice with PBS. Immunoprecipitated proteins were eluted from the support in Laemlli loading buffer, separated by SDS-PAGE and detected by Western blotting using ColXVIII (QH48.14) *(54)*, EGFR (Cell Signaling Technology #4267), HER2 (Cell Signaling Technology #4290), integrin α6 (BD Biosiences #555734) and integrin β1 (Sigma Aldrich #AB1952) antibodies (Table S2).

**Fluorescence-activated cell sorting (FACS)**

The single cell suspensions from human BC cell lines were prepared by trypsinization, after which the action of trypsin was blocked with complete medium containing FBS, centrifuged and resuspended in PBS for further analyses. Mouse mammary tumor tissue was extracted, minced and incubated in RPMI-1640 (Sigma-Aldrich) medium containing 0.3% collagenase, type 1 (CLS-1, Worthington, USA) for 1 hour at 37°C, and filtered through a 70-µm mesh (BD Biosciences) to obtain single cell suspensions. The cell suspensions were then washed with PBS containing 4% FBS and taken for FACS analyses. All the samples prepared from mouse tissues or human cell lines were analysed on either LSR Fortessa or Aria II flow cytometers equipped with FACSDiva Software, or on a FACS Calibur cytometer equipped with CellQuest software (all from BD Biosciences). The following antibodies were used: fluorescent anti-CD24, -CD29, -CD44, -CD49f and Lineage antibody cocktail (CD3, CD11b, CD45R/B220, Ter-119, Ly-6G/C) (Table S3). Enriched mammary SC or putative CSC populations from mouse mammary tumors were immunophenotyped as ($Lin^{-}CD24^{+}CD44^{+}CD29^{High}$ $CD49f^{High}$) and those from human cell lines as ($CD44^{+}CD24^{-/low}$). All FACS data were analysed using Flowjo software (version 7.6.3, Tree Star Inc.).

**Colony-forming unit assay**

The MCF7 cells were sorted with a BD Aria II flow cytometer for breast CSC ($CD44^{+}CD24^{-/low}$) and non-breast CSC (nCSC) ($CD44^{+}CD24^{+/Hi}$) cells. For this purpose the cells were suspended in a 1:1 mixture of ice-cold DMEM and growth factor-reduced Matrigel matrix (BD Biosciences, #354230) and plated in a 6-well plate. The gels were kept at +37 ° C for 15 min to allow them to solidify, and DMEM was added on top of the matrix. The formation of spherical 3D colonies was analysed after 7-10 days of plating. The colonies were imaged by phase contrast microscopy and CSC characteristics were defined based on the spheroid morphology.

**ErbB inhibitor treatments in human breast cancer cell lines**

Human BC that expressed high levels of HER2 receptor (SKBR-3, JIMT-1 and BT474) and triple negative MDA-MB-231 were used for the drug tests. Cells were seeded at a density of $1{\times}10^4$ cells per well in a 96-well format. For *COL18A1* gene expression knockdown, the two siRNAs that target ColXVIII, or scrambled siRNA as a negative control, were transfected into the cells as explained above. 48 hours post transfection the cells were washed with growth medium and EGFR/HER2 targeting drugs were added (lapatinib in DMSO at a final concentration of 2µM), trastuzumab (at 12.5µg/ml) or panitumumab (at 1.5µg/ml). The control scrambled and control ColXVIII KD cells received the vehicle only instead of the drugs. Cell proliferation and migration were monitored for 30-120 h after drug treatment using the IncuCyte live cell analysis system as described above in connection with the cell proliferation and migration assays.

#### *In vivo* lapatinib treatment

WT-PyMT and $18^{-/-}$-PyMT mice were used for the *in vivo* lapatinib efficacy study. 7-weeks old female mice were treated with two different doses of lapatinib ditosylate (LC laboratories), either 35 or 70mg per kilogram of body weight (mpk) twice a day and five days a week for a total of three weeks. The dose was chosen to be half the optimized human dose according published guidelines *(61)*. Lapatinib was given as an oral suspension in a vehicle consisting of water with 0.5% hydroxypropyl methylcellulose and 0.1% Tween-80. Vehicle-treated WT-PyMT and $18^{-/-}$-PyMT mice were included as controls. The animals were regularly monitored with respect to body weight and tumor development. The mammary glands were collected after three weeks of treatment, weighed at the time of sacrifice and samples taken for processing and further analyses.

### Supplementary Figures

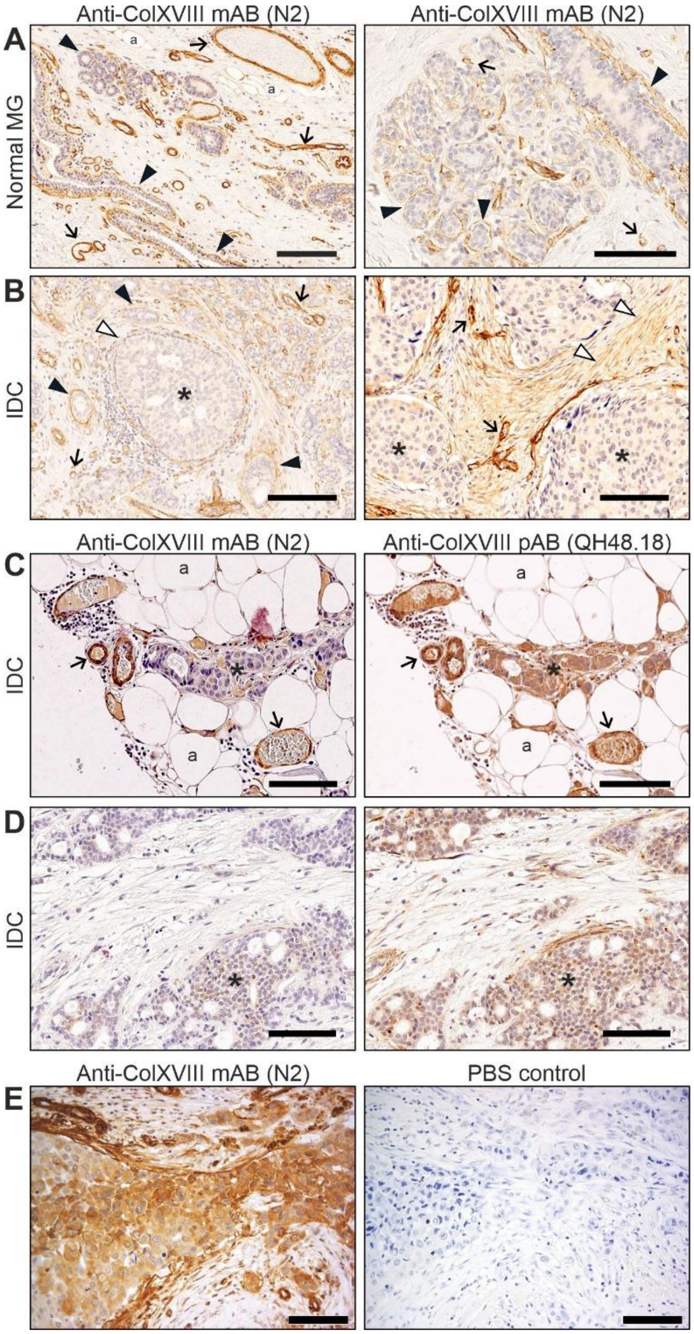

**Fig. S1. ColXVIII immunohistochemistry in human breast cancer and the normal mammary gland. (A-B)** A monoclonal anti-ColXVIII antibody (mAB) DB144-N2 *(53)* was used to detect ColXVIII in human breast tissues. **(A)** Normal mammary gland (MG)**, (B)** invasive ductal carcinoma (IDC). **(C-D)** Comparison of ColXVIII signals in sequential IDC sections stained with two anti-ColXVIII antibodies, the monoclonal DB144-N2 and the polyclonal QH48.18 antibody (pAB) *(54)*. **(E)** DB144-N2 staining of HER2 type IDC on the left (also shown in the main Fig. 1G) and its negative control in which the primary antibody was replaced with PBS (right). Black arrowhead, basement membrane (BM); white arrowhead, ColXVIII absent from the BM; arrow, vascular BM; asterisk, cytoplasmic ColXVIII staining in tumor cells; a, adipocyte. Scale bars 100µm.

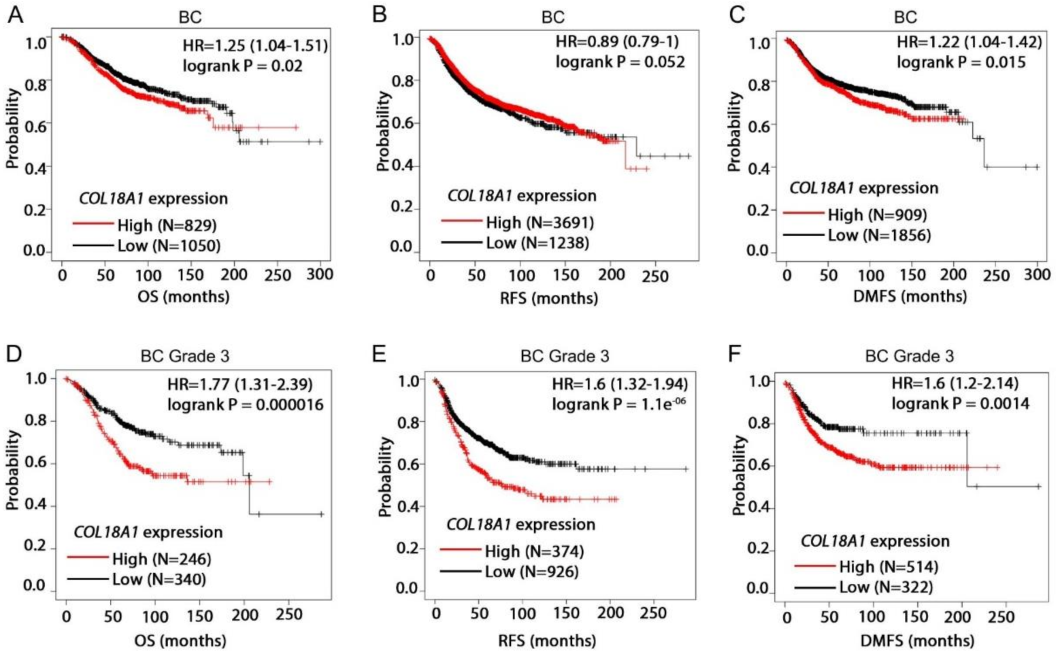

**Fig. S2. Association between ColXVIII expression and breast cancer patient survival.** Kaplan-Meir plots showing overall survival (OS), relapse-free survival (RFS) and distant metastasis-free survival (DMFS) of BC patients stratified by ColXVIII expression levels (probe: 209082_s_at). **(A-C)** All available BC cases and **(D-F)** grade 3 BC cases. The open access gene expression data and patient survival information from TCGA, GEO and EGA compiled in a single database at www.kmplot.com *(24)* were used for the meta-analyses. High ColXVIII, red line; low ColXVIII, black line. Hazard ratio (HR) and log-rank P values were automatically computed using the best-performing threshold as the cutoff. The number of patients for each analysis is indicated in brackets on the respective survival graphs.

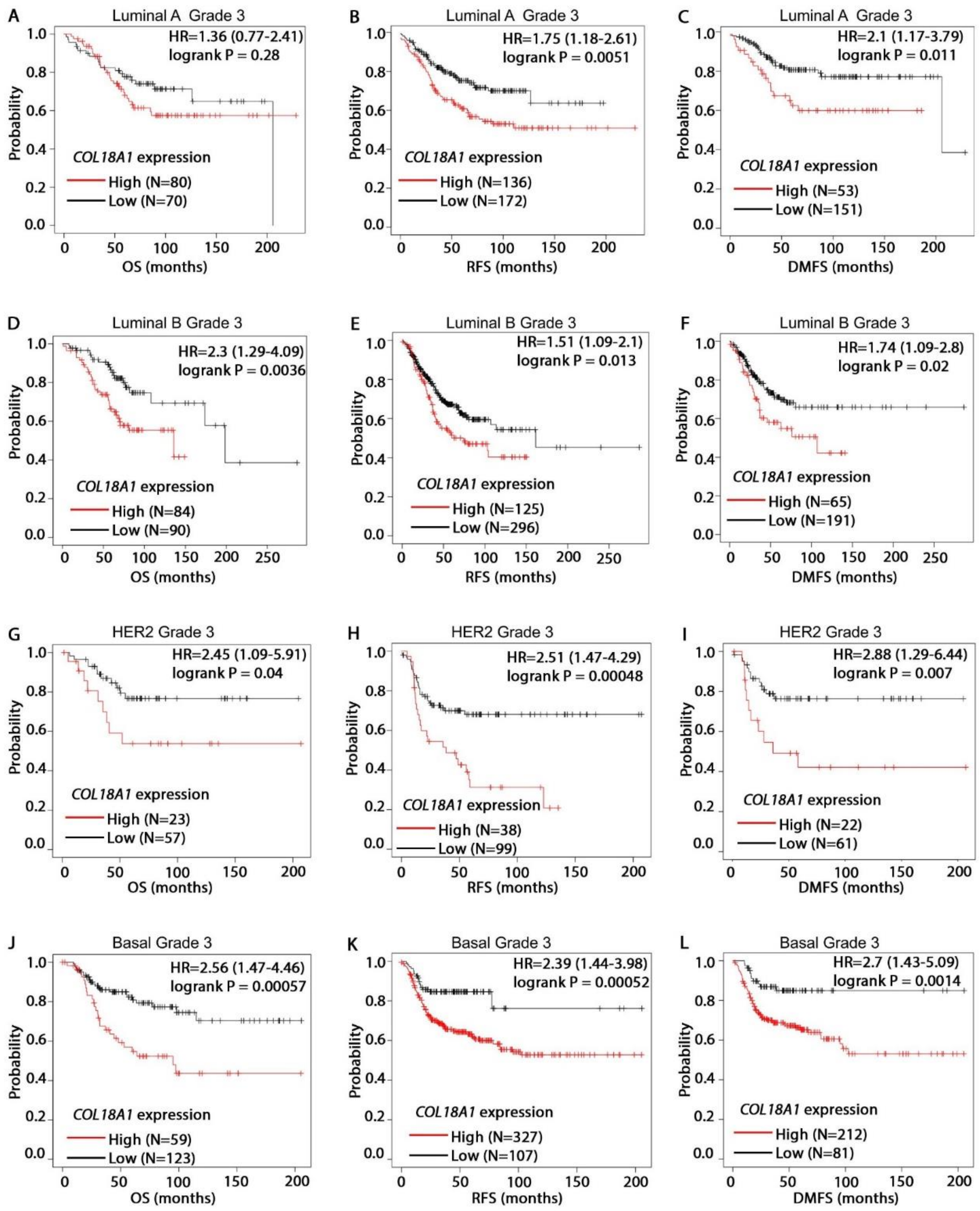

**Fig. S3. ColXVIII expression and patient survival in grade 3 breast tumors of different molecular subtypes.** Kaplan-Meir plots showing overall survival (OS), relapse-free survival (RFS) and distant metastasis-free survival (DMFS) of BC patients stratified by ColXVIII expression levels (probe: 209082_s_at). **(A-C)** Luminal A, **(D-F)** luminal B, **(G-I)** HER2 and **(J-L)** basal subtype of grade 3 BCs. The open access gene expression data and patient survival information from TCGA, GEO and EGA compiled in a single database at www.kmplot.com *(24)* were used for the meta-analyses. High ColXVIII, red line; low ColXVIII, black line. Hazard ratio (HR) and log-rank P values were automatically computed using the best-performing threshold as the cutoff. The number of patients for each analysis is indicated in brackets on the respective survival graphs.

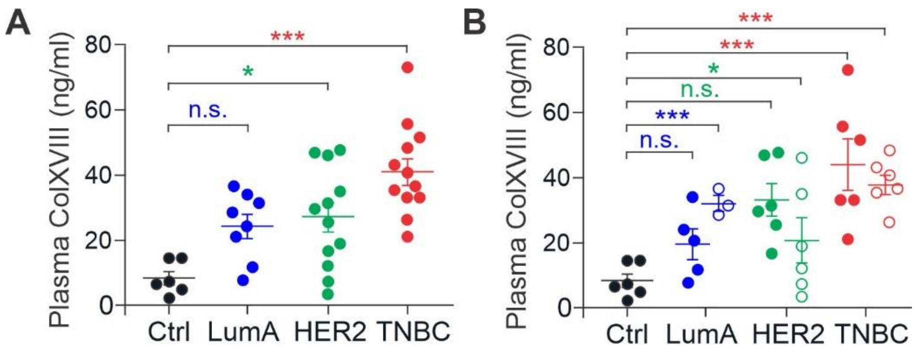

**Fig. S4. ColXVIII in breast cancer patient plasma. (A)** Levels of circulating ColXVIII (ng/ml) in plasma samples from patients with luminal A (N=8), HER2 (N=12) and basal (N=12) types of BC by comparison with healthy controls (N=6). **(B)** Levels of circulating ColXVIII (ng/ml) in plasma samples from BC patients classified according to the molecular subtype, and with and without lymph node metastasis. Numbers of samples: node negative N=5, node positive N=3 for luminal A-type BC; node negative N=6, node positive N=6 for the HER2 type; and node positive N=6, node negative N=6 for the basal/TNBC type. ColXVIII concentrations were measured using a monoclonal N-terminal TSP-1 domain-targeting anti-ColXVIII antibody (DB144-N2) *(53)* in an indirect ELISA assay. Student's 't' test was used for statistical analysis. *, $p<0.05$. ***, $p<0.001$; and n.s., not significant. Error bars represent s.e.m.

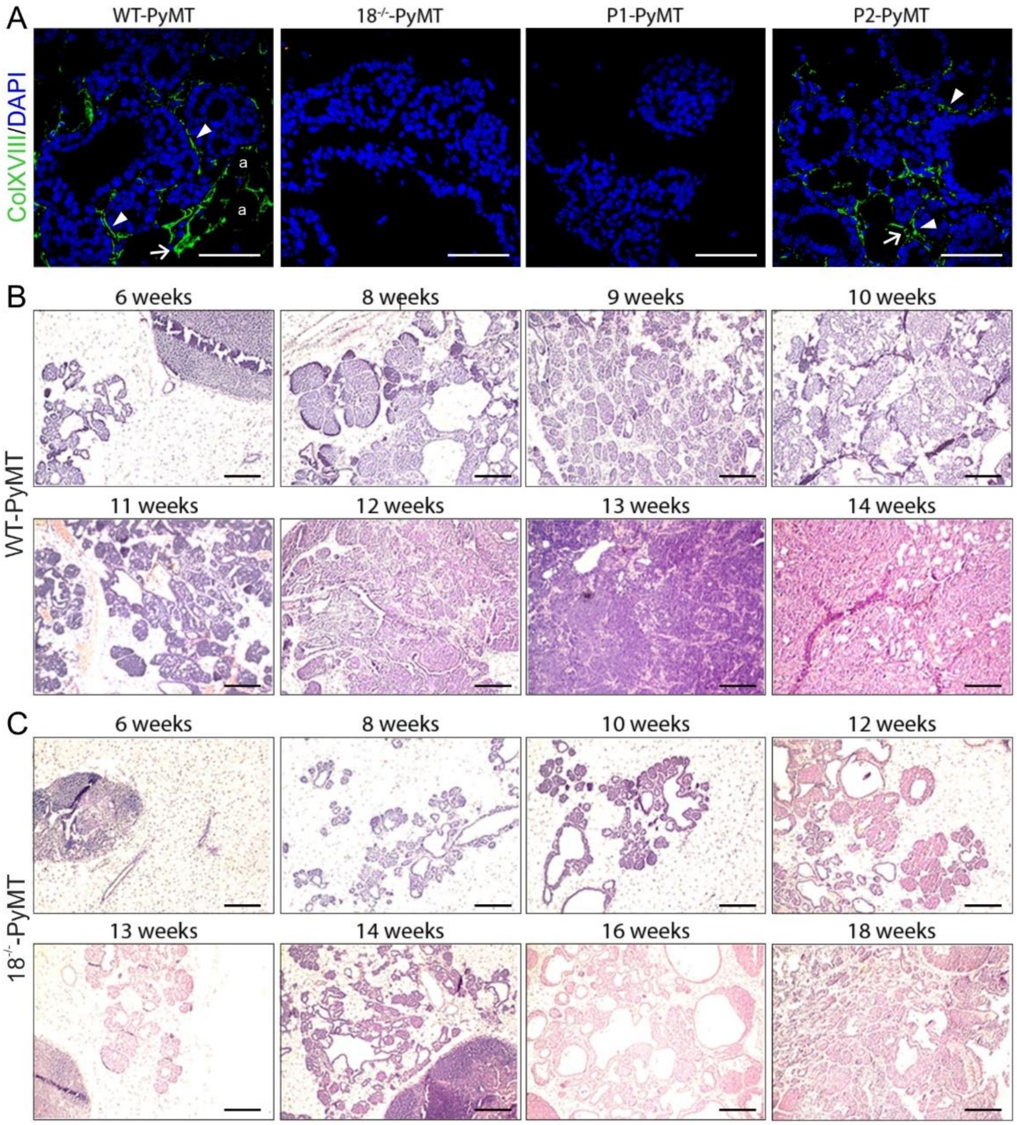

**Fig. S5. The short ColXVIII isoform supports mammary carcinogenesis in the MMTV-PyMT model. (A)** Immunofluorescence staining of ColXVIII (green) in wild type MMTV-PyMT (WT-PyMT) mammary tumors, in $18^{-/-}$-PyMT tumors lacking all ColXVIII isoforms, in P1-PyMT tumors lacking the short ColXVIII and expressing long/medium ColXVIII, and in P2-PyMT tumors lacking the long/medium isoforms and expressing exclusively the short ColXVIII. The staining shows that the short ColXVIII is the prevalent isoform in mammary tumors. A polyclonal antibody targeting the N-terminal TSP-1 domain of mouse ColXVIII was used in the staining *(59)*. Scale bars, 100μm. Arrowhead, ductal basement membrane; arrow, blood vessel; a, adipocyte. **(B-C)** Hematoxylin-eosin staining of mammary tumors harvested from the WT-PyMT mice at the age of 6-14 weeks and from $18^{-/-}$-PyMT mice at age 6-18 weeks. The mammary tumors transformed to carcinomas between weeks 10 and 13 in the WT-PyMT mice, but between weeks 16 and 18 in the $18^{-/-}$-PyMT mice. The tumors were classified according to *(62)*. Scale bars, 200μM.

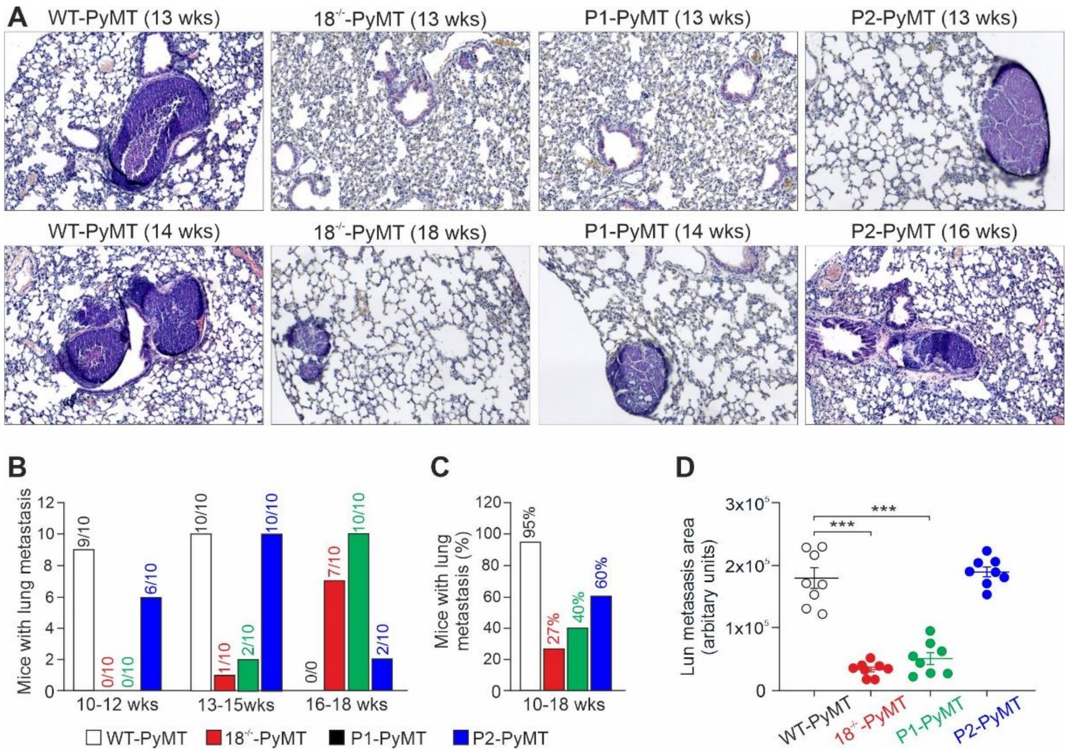

**Fig. S6. Lung metastasis in wild type MMTV-PyMT and ColXVIII-deficient PyMT mice. (A)** Hematoxylin-eosin-stained lung sections of WT-PyMT, $18^{-/-}$-PyMT, P1-PyMT and P2-PyMT mice. **(B)** Numbers of mice with one or more lung metastases in the four mouse lines by age groups. The total number of mice of each genotype was ten, except for the WT-PyMT mice, which could not be monitored beyond 14 weeks for ethical reasons. **(C)** Percentages of mice with lung metastasis in the PyMT mouse lines at weeks 10-18. **(D)** ImageJ analysis of tumor areas in lung samples at weeks 13-18. One lung section per mouse was analysed, and N=8 for all genotypes. Student's 't' test was used for the statistical analysis. ***, p<0.001; n.s., not significant.

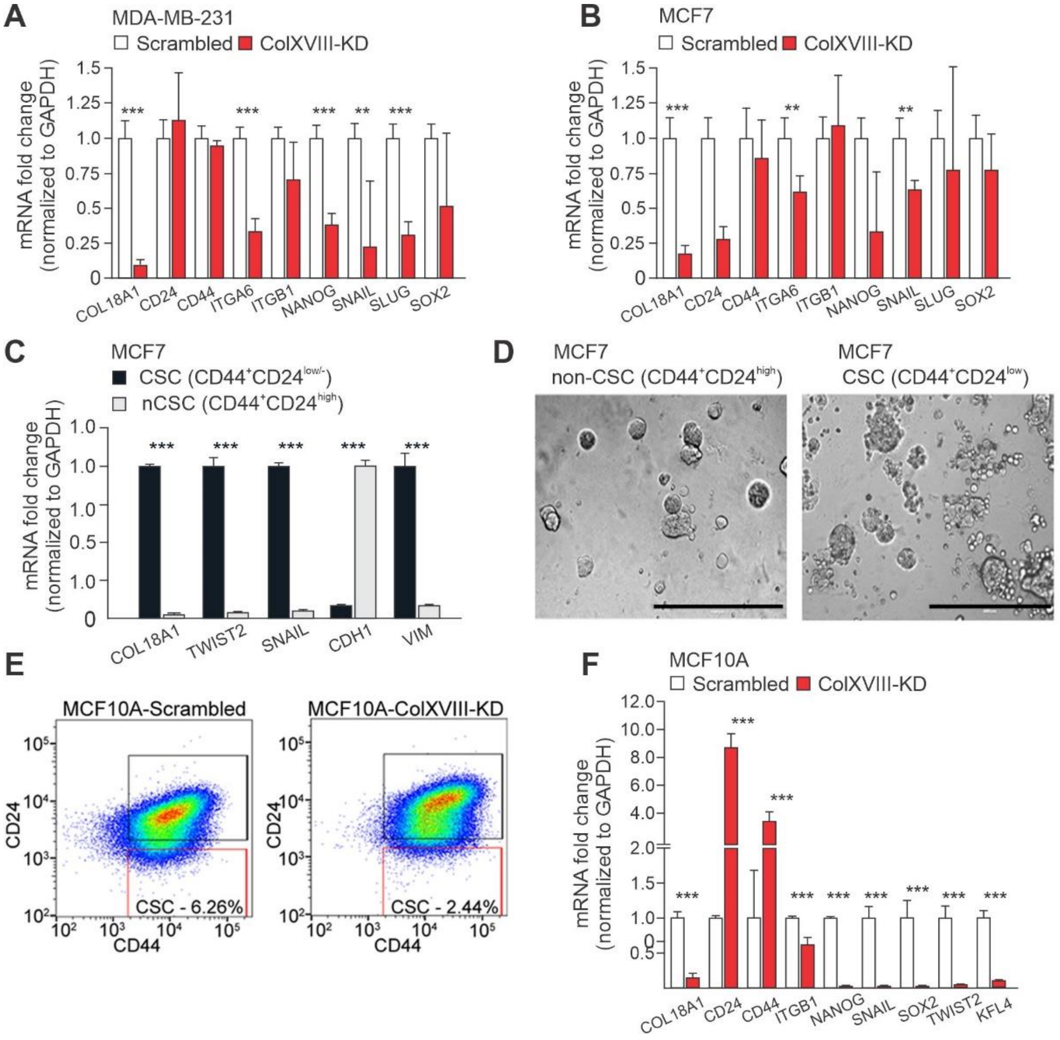

**Fig. S7. ColXVIII supports breast cancer stem cells. (A-B)** Quantitative RT-PCR analysis of relative ColXVIII expression levels (normalized to GAPDH) and stem cell marker gene transcripts in siRNA-based ColXVIII knockdown (KD) and scrambled vector-transfected MDA-MB-231 and MCF7 BC cell lines. **(C)** Quantitative RT-PCR analysis of relative ColXVIII expression levels (normalized to GAPDH) and stem cell marker gene transcripts in human BC stem cells (CSC) ($CD44^{+}/CD24^{low/-}$) and non-CSC ($CD44^{+}/CD24^{high}$) populations sorted by flow cytometry from MCF7 cells. **(D)** Colony-forming unit assay of CSC and non-CSC populations of MCF7 cells cultured in growth factor-reduced Matrigel. Scale bar 500µm. **(E)** CSC populations ($CD44^{+}/CD24^{low/-}$) in the ColXVIII-KD and scrambled vector-transfected MCF10A cells analysed by flow cytometry. **(F)** Quantitative RT-PCR of relative ColXVIII expression levels (normalized to GAPDH) and stem cell marker gene transcripts in ColXVIII-KD and scrambled MCF10A cells. Student's 't' test was used for the statistical analysis. **, $p<0.01$; ***, $p<0.001$. Error bars represent s.e.m.

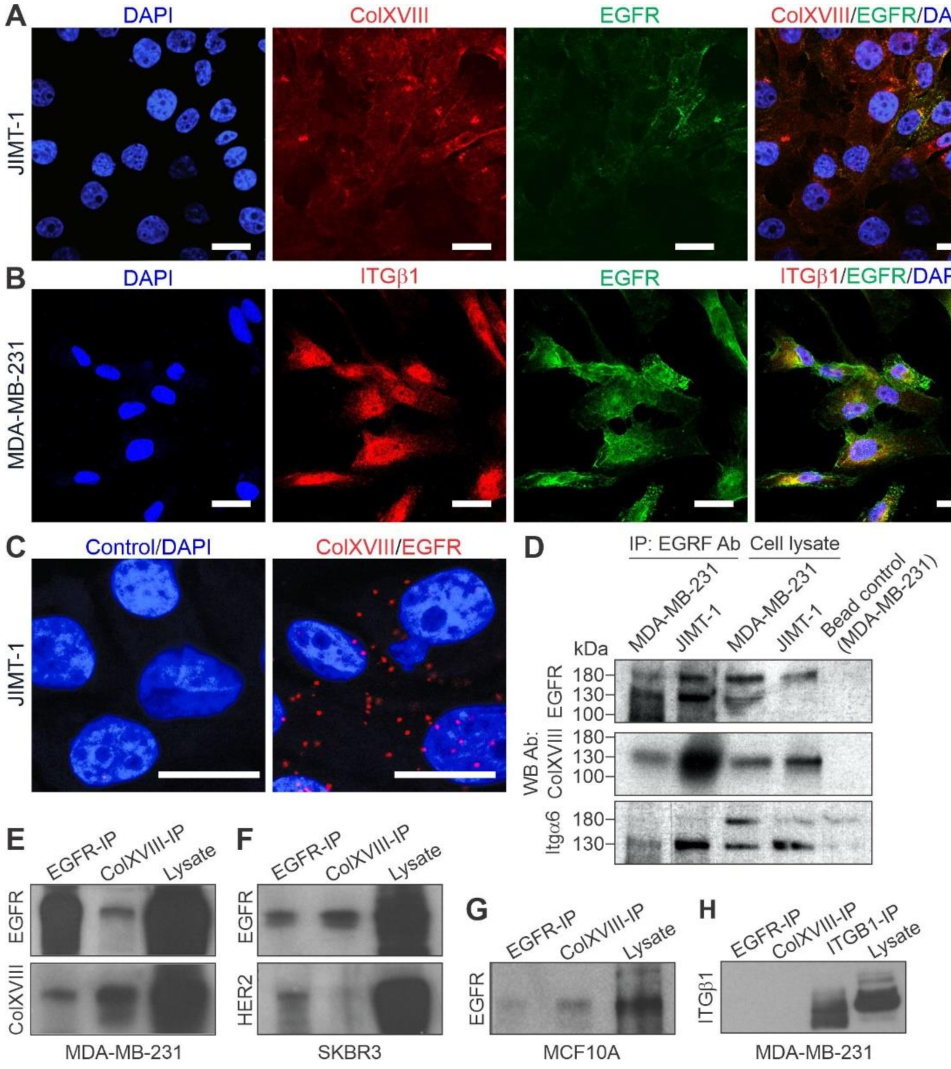

**Fig. S8. Interactions between ColXVIII, ErbBs and integrins.** Representative images of immunofluorescence staining of **(A)** ColXVIII (red) and EGFR (green) in JIMT-1 cells, and **(B)** integrin β1 (red) and EGFR (green) in MDA-MB-231 cells. The nuclei are counterstained with DAPI. Scale bars, 20μm. **(C)** *In situ* proximity ligation assay. Evidence of proximity (distance less than 40 nm) for ColXVIII (DB144-N2[29]) and EGFR (Abcam #52894) in JIMT-1 cells is indicated by the presence of red dots. The negative control without primary antibodies is also shown. Scale bars, 20μm **(D)** Co-immunoprecipitation (Co-IP) assay with an EGFR antibody (Abcam #52894) in HER2-negative MDA-MB-231 cells and HER2-positive JIMT-1 cells. Protein complexes were detected by Western blotting (WB) with anti-EGFR (Cell Signaling Technology, #4267), anti-ColXVIII (QH48.14) *(54)* and anti-α6 integrin (Abcam, #ab97760) antibodies. A goat anti-mouse IgG coated magnetic bead control is shown in the MDA-MB-231 cells. **(E-H)** Co-IP with EGFR (#52894) and ColXVIII (DB144-N2) antibodies in MDA-MB-231 **(E)**, SKBR3 **(F)** and MCF10A **(G)** cell lysates, and with EGFR, ColXVIII (QH48.14) and integrin β1 (Millipore AB1952) antibodies in MDA-MB-231 lysates. Protein complexes were detected by WB with the EGFR (Cell Signaling Technology, #4267), ColXVIII (QH48.14), HER2 (Cell Signaling Technology, #4290) and integrin β1 (pAB, Millipore-AB1952) antibodies (Table S2).

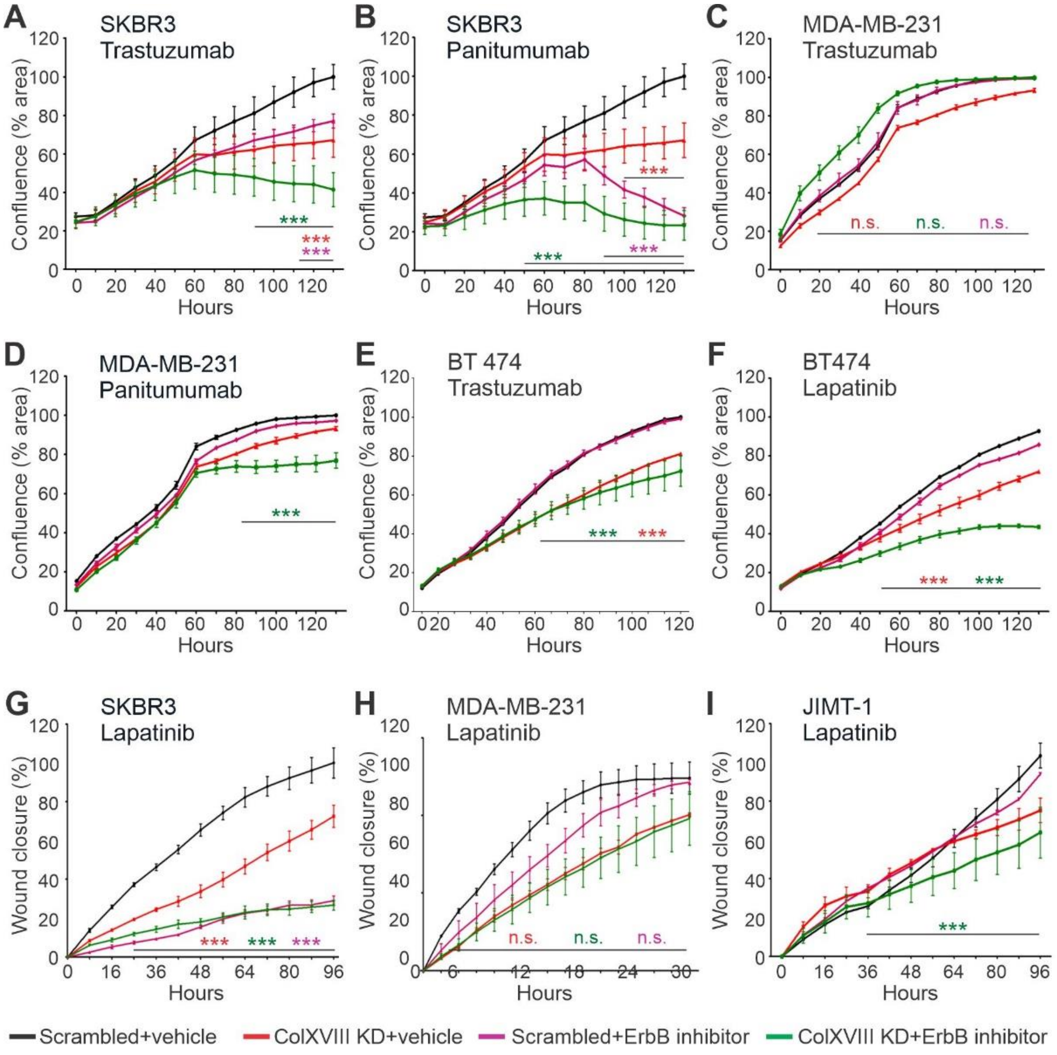

**Fig. S9. Analysis of breast cancer cell proliferation and migration upon ColXVIII knockdown and/or EGFR/ErbB-targeting drug treatment. (A, B)** siRNA-based knockdown (KD) of ColXVIII expression significantly improves the efficacy of the HER2-targeting humanized monoclonal antibody (mAB) trastuzumab and the EGFR-targeting humanized mAB panitumumab in HER2-amplified SKBR3 cells. **(C)** Trastuzumab is not effective in triple-negative MDA-MB-231 cells, whereas **(D)** panitumumab slowed down the proliferation of these cells upon ColXVIII-KD. **(E-F)** The luminal B type, HER2-expressing BC cell line BT474 has an activating mutation in the PI3K signal mediator and is thus resistant to ErbB2-targeting drugs, but ColXVIII-KD can dismantle resistance to lapatinib. **(G-I)** Both ColXVIII siRNAs and lapatinib alone lead to reduced migration of SKBR3 cells, and to reduced migration of JIMT-1 cells when administered simultaneously, whereas MDA-MB-231 cell migration is not affected by either treatment. Student's 't' test was used for the statistical analysis. ***, $p<0.001$; n.s., not significant. Error bars represent s.e.m.

### Supplementary Tables

#### Table S1. Human breast cancer samples with histopathological and relapse data

| | **Umeå cohort** (N=95) | | **Oulu cohort** (N=21) | |
|---|---|---|---|---|
| | number | percentage | number | percentage |
| **Oestrogen (ER) receptor** | | | | |
| Positive | 75 | 79 % | 11 | 52 % |
| Negative | 18 | 19 % | 10 | 48 % |
| NA | 2 | 2 % | | |
| **Progesterone (PR) receptor** | | | | |
| Positive | 55 | 58 % | 10 | 48 % |
| Negative | 38 | 40 % | 11 | 52 % |
| NA | 2 | 2 % | | |
| **HER2 receptor** | | | | |
| Positive | 20 | 21 % | 13 | 62 % |
| Negative | 73 | 77 % | 8 | 38 % |
| NA | 2 | 2 % | | |
| **Nuclear grade** | 18 | 19 % | 2 | 10 % |
| 1 (low) | 63 | 66 % | 4 | 19 % |
| 2 (moderate) | 14 | 15 % | 15 | 71 % |
| 3 (high) | | | | |
| **Ki67 status** | 47 | 50 % | 4 | 19 % |
| Low | 42 | 44 % | 16 | 76 % |
| High | 6 | 6 % | 1 | 5 % |
| NA | | | | |
| **Relapse** | | | | |
| Yes | 25 | 26% | NA | NA |
| No | 70 | 74% | NA | NA |

NA, data not available

### Table S2. Antibodies

| Target/antibody | Catalog # | Manufacturer/Reference |
|---|---|---|
| Cy3-conjugate αSMA | C6198 | Sigma |
| Cleaved Caspase-3 | MAB835 | R&D Systems |
| Collagen XVIII (human) | QH4818 | In House *(54)* |
| Collagen XVIII (human) | DB144-N2 | In House *(53)* |
| Collagen XVIII (mouse) | anti-NC11 | In House *(59)* |
| Cyclophyllin B | SAB4200201 | Sigma Aldrich |
| Cytokeratin 5 | PRB-160P | Covance |
| EGFR (D38B1) | 4267 | Cell Signaling Technology |
| EGFR | AB2430 | Abcam |
| EGFR | AB52894 | Abcam |
| FITC-conjugated CD44 | 103005 | Biosite |
| HER2/ErbB2 (D8F12) | 4290 | Cell Signaling Technology |
| Integrin alpha 6 (human) | ab97760 | Abcam |
| Integrin alpha 6 (human) | 555734 | BD Biosciences |
| Integrin alpha 6 (mouse) | MAB1356 | Chemicon |
| Integrin beta 1 (human) | AB1952 | Sigma Aldrich |
| Integrin beta 1 (human) | SC9970 | Santa Cruz |
| Ki67 | M7248 | DakoCytomation |
| Lineage antibody cocktail (CD3, CD11b, CD45R/B220, Ter-119, Ly-6G/C) | 558074 | BD Biosciences |
| pan44/42 MAPK (Erk1/2) (137F5) | 4695S | Cell Signaling Technology |
| phospho AKT (Tyr-326) | sc-109904 | Santa Cruz |
| panAKT (c67e7) | 4691 | Cell Signaling Technology |
| PE -conjugated CD29 | 102207 | Biolegend |
| PE/Cy7-conjugated CD24 | 101821 | Biolegend |
| PE -conjugated CD49f | 12-0495-82 | eBiosciences |
| Phospho-Akt (Ser473) (D9E) XP | 4060S | CST |
| Phospho-EGFR (Tyr1068) (D7A5) | 3777 | CST |
| Phospho-HER2/ErbB2 (Tyr1221/1222) (6B12) | 2243 | CST |
| Phospho-p38 MAPK (Thr180/Tyr182) (12F8) | 4631S | CST |
| Phospho-p44/42 MAPK (Erk1/2) (Thr202/Tyr204) | 4370S | CST |
| Phospho-PI3K p85 (Tyr458)/p55 (Tyr199) | 4228 | CST |
| PI3 Kinase Class III (D4E2) | 3358 | CST |
| PI3 Kinase p110α (C73F8) | 4249 | CST |
| PI3 Kinase p110γ (D55D5) | 5405 | CST |
| PI3 Kinase p110ß (C33D4) | 3011 | CST |
| PI3 Kinase p85 (19H8) | 4257 | CST |
| Tubulin (Beta) | MAB-3408 | Millipore |

**Table S3.** qRT-PCR Primers

| Gene Symbol | Gene Name | Primers | Sequences (5' → 3') |
|---|---|---|---|
| *COL18A1* | Collagen, type XVIII, alpha 1 chain (C-terminal, endostatin, all isoforms) | Forward | TGCCCATCGTCAACCTCAAG |
| | | Reverse | CAGAGCCTGAGAACAGAGCC |
| *COL18A1* | Collagen, type XVIII, alpha 1 chain (N-terminal, Short isoform) | Forward | CTCCTGGACGTGCTCGC |
| | | Reverse | TCATCCGTCTGGGTGACCT |
| *COL18A1* | Collagen, type XVIII, alpha 1 chain (N-terminal, Medium isoform) | Forward | GCCGTGGCATTCCTAGCTC |
| | | Reverse | CTGATGCGCTCTGAAGATGGT |
| *COL18A1* | Collagen, type XVIII, alpha 1 chain (N-terminal, Long isoform) | Forward | AGCTTCTCTCTCCTCCTTGCT |
| | | Reverse | GCGAGAGTCCTTGGCTGTCT |
| *Col18a1* | Collagen, type XVIII, alpha 1 chain (C-terminal, endostatin, all isoforms) | Forward | GTGCCCATCGTCAACCTGAA |
| | | Reverse | GACATCTCTGCCGTCAAAAGAA |
| *Col18a1* | Collagen, type XVIII, alpha 1 chain (N-terminal, Short isoform) | Forward | GGATGTGCTCACCAGTTTGG |
| | | Reverse | GTCATCGATTTGTGAGATCTTC |
| *Col18a1* | Collagen, type XVIII, alpha 1 chain (N-terminal, Medium isoform | Forward | ACCTCCAGGCACCACTGGGA |
| | | Reverse | GCCCGACGTGAGGGTCATCG |
| *Col18a1* | Collagen, type XVIII, alpha 1 chain (N-terminal, Long isoform) | Forward | AGGAGGACGGGTACTGTGTG |
| | | Reverse | TGAGGGTCATCGATTTGTGA |
| *CD24* | CD24 molecule | Forward | CTCCTACCCACGCAGATTTATTC |
| | | Reverse | AGAGTGAGACCACGAAGAGAC |
| *CD44* | CD44 molecule | Forward | CTGCCGCTTTGCAGGTGTA |
| | | Reverse | CATTGTGGGCAAGGTGCTATT |
| *CDH1* | Cadherin 1 | Forward | ATTTTTCCCTCGACACCCGAT |
| | | Reverse | TCCCAGGCGTAGACCAAGA |
| *GAPDH* | glyceraldehyde-3-phosphate dehydrogenase | Forward | ACAACTTTGGTATCGTGGAAGG |
| | | Reverse | GCCATCACGCCACAGTTTC |
| *ITGA6* | Integrin alpha 6 | Forward | GGCGGTGTTATGTCCTGAGTC |
| | | Reverse | AATCGCCCATCACAAAAGCTC |
| *ITGB1* | Integrin beta 1 | Forward | GTAACCAACCGTAGCAAAGGA |
| | | Reverse | TCCCCTGATCTTAATCGCAAAAC |
| *KLF4* | Kruppel like factor 4 | Forward | CAGCTTCACCTATCCGATCCG |
| | | Reverse | GACTCCCTGCCATAGAGGAGG |
| *NANOG* | Nanog homeobox | Forward | CCCCAGCCTTTACTCTTCCTA |
| | | Reverse | CCAGGTTGAATTGTTCCAGGTC |
| *SNAIL1* | Snail family zinc finger 1 | Forward | GTGTCTCCCAGAATATT |
| | | Reverse | GTTTGAAATATAAATACCAGTGT |
| *SOX2* | SRY-box 2 | Forward | TACAGCATGTCCTACTCGCAG |
| | | Reverse | GAGGAAGAGGTAACCACAGGG |

**Table S3.** qRT-PCR Primers

| Gene Symbol | Gene Name | Primers | Sequences (5' → 3') |
|---|---|---|---|
| *SLUG* | Snail family transcriptional repressor 2 | Forward | GCTGATGGCTAGATTGAG |
| | | Reverse | ATCCTATTACAGACTCTATTACAA |
| *TWIST2* | Twist basic helix-loop-helix transcription factor 2 | Forward | TTCTCCGTGATTGCTTGG |
| | | Reverse | AGGATACACAGCCACACT |
| *VIM* | Vimentin | Forward | AGTCCACTGAGTACCGGAGAC |
| | | Reverse | CATTTCACGCATCTGGCGTTC |
